# Truncating *RELA* variants drive autoinflammation and autoimmunity by impairing the negative feedback control of NF-kB

**DOI:** 10.1101/2025.11.09.687461

**Authors:** Nadja Lucas, Sophia Weidler, Antonia Eicher, Baerbel Keller, Özlem Satirer, Adam Desrochers, Timothy Ramnarine, Mohammad Mokhtari, Sophie Elstner, Timmy Strauss, Simon Mages, Arek Kendirli, Oana Buzoianu, Lina Igel, Tim Niehues, Sandra von Hardenberg, Maria Fasshauer, Rami Abou Jamra, Hagen Ott, Ulrike Hüffmeier, Catharina Schütz, Marisa Bijwaard, Susan Wagner, Paulina Switala, Sarah Koss, Jurek Schultz, Stefanie Kretschmer, Jasmin Kümmerle-Deschner, Christine Wolf, Johanna Klughammer, Min Ae Lee-Kirsch

## Abstract

The NF-κB signaling pathway coordinates inflammation, cell survival, and proliferation, while restraining excessive cell death to maintain immune homeostasis. Truncating mutations in *RELA*, encoding the NF-κB subunit p65, have been linked to autoinflammation and autoimmunity, but the underlying mechanisms remain incompletely defined. We investigated six patients from five unrelated families carrying novel heterozygous truncating *RELA* variants. Despite reduced p65 expression, patients exhibited a broad spectrum of inflammatory manifestations alongside elevated baseline and stimulus-induced pro-inflammatory cytokines. Functional analyses in patient-derived cells and mutant *RELA* knock-in models showed that upstream NF-κB signaling was intact, but induction of inhibitory regulators such as IκBα and A20 was impaired. This defective feedback control shifted immune homeostasis toward amplified inflammatory responses that depended on the residual activity of the remaining functional *RELA* allele. Single-cell transcriptomics revealed distinct cell type-specific consequences: monocytes displayed constitutive type I interferon and NF-κB activation, B cells retained partial compensatory signaling, whereas T and NK cells exhibited transcriptional signatures of cell death pathways. Patient fibroblasts and mutant *RELA* knock-in cells further confirmed enhanced TNF-induced inflammatory gene expression and hypersensitivity to apoptosis and necroptosis. These findings establish *RELA* haploinsufficiency as a cause of systemic immune dysregulation, and link defective NF-κB feedback control to unchecked inflammation and inflammatory cell death.

**Graphical abstract:** 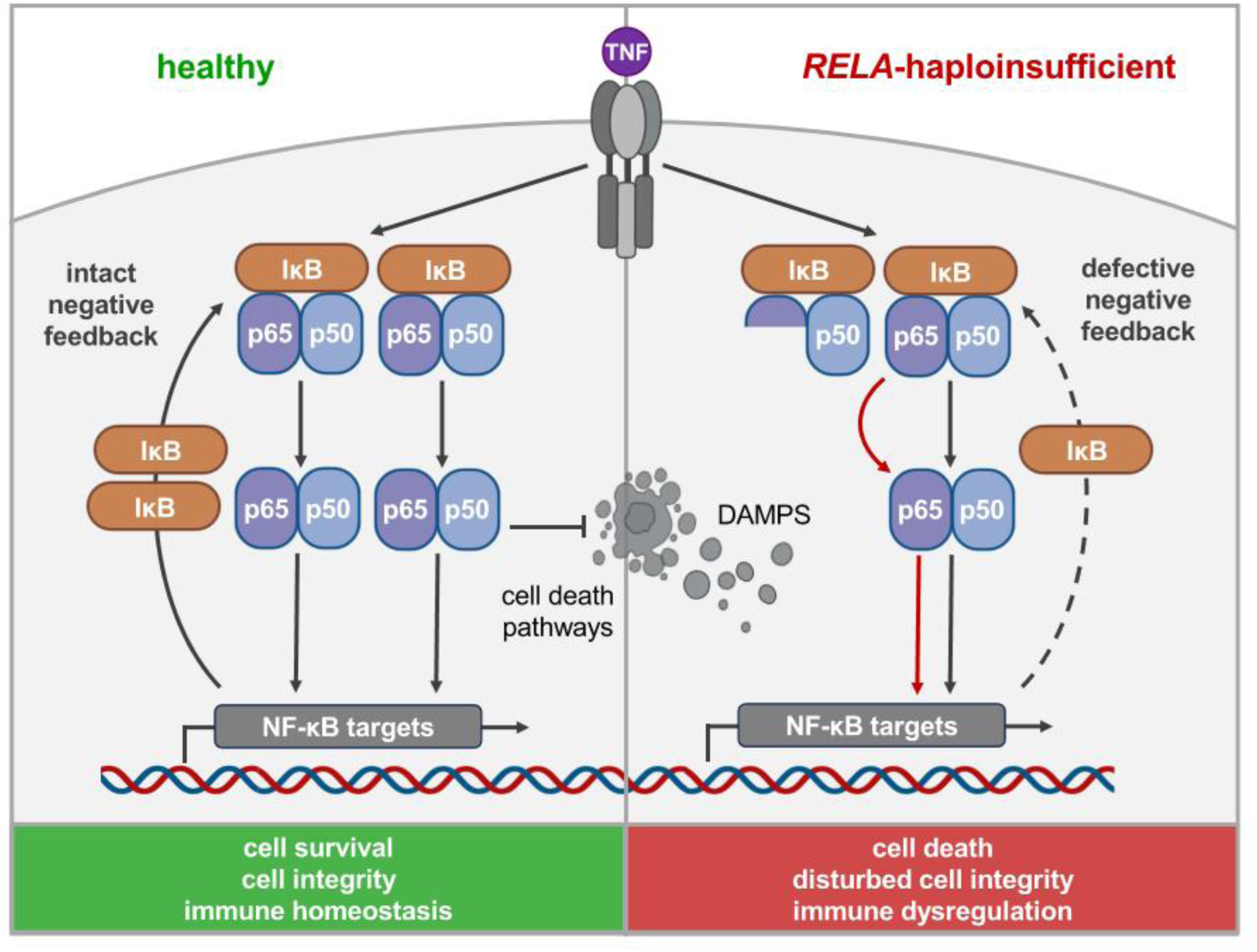

## Introduction

The nuclear factor kappa-light-chain-enhancer of activated B cells (NF-κB) signaling pathway is a complex regulatory network that plays a central role in immune responses, inflammation, and cell survival (1–3). It functions primarily as a transcriptional activator of genes involved in both innate and adaptive immunity, and is critical for maintaining immune homeostasis and responding to cellular stress. The NF-κB family consists of five structurally related proteins: RELA (p65), RELB, c-Rel, NF-κB1 (p105/p50), and NF-κB2 (p100/p52) (1). All subunits share a conserved Rel homology domain (RHD) responsible for dimerization, DNA binding, and interaction with inhibitory IκB proteins. Among these, RELA, RELB, and c-Rel possess C-terminal transactivation domains (TADs) and act as transcriptional activators (1). In contrast, p50 and p52 lack TADs and must form heterodimers with TAD-containing subunits to activate gene transcription. In the absence of stimulation, NF-κB dimers are sequestered in the cytoplasm through interactions with inhibitors of κB (IκBs), such as IκBα, IκBβ, and IκBε, which mask their nuclear localization sequences (NLS), thereby preventing nuclear translocation (1, 2). The canonical NF-κB pathway is activated by a wide range of stimuli, including tumor necrosis factor (TNF), interleukin-1 (IL-1), TLR ligands, and antigens (1, 2).

Upon activation of these receptors, upstream kinases converge on the IκB kinase (IKK) complex, composed of IKKα, IKKβ, and the regulatory subunit NEMO (IKKγ). The activated IKK complex phosphorylates IκB proteins, marking them for ubiquitination and proteasomal degradation (1, 2). This process releases the NF-κB dimers, most commonly p65/p50, which translocate into the nucleus, bind κB enhancer elements in target gene promoters, and induce transcription of a broad array of genes involved in inflammation, cell proliferation, survival, and immune regulation (1, 2).

The non-canonical NF-κB pathway, activated by CD40, BAFFR, or LTβR, relies on NIK-driven processing of p100 to p52, which dimerizes with RelB to form the active complex (3). Both pathways are tightly regulated by negative feedback mechanisms. In the canonical pathway, this includes NF-κB-induced expression of IκBα and deubiquitinases such as TNFAIP3 (A20), CYLD, and OTULIN, which act to terminate upstream signaling cascades (4). Dysregulation of NF-κB signaling is associated with a range of pathologies, including chronic inflammation, autoimmune diseases, and cancer (1, 2).

Truncating *RELA* variants have been implicated in a spectrum of autoinflammatory and autoimmune disorders, including Behçet-like mucocutaneous ulcerations (5–9), inflammatory bowel disease (10, 11), juvenile idiopathic arthritis (11), autoimmune hematological conditions (12), and early-onset systemic lupus erythematosus (13). While both haploinsufficiency, leading to increased sensitivity to TNF, and dominant-negative effects, associated with heightened TLR7 signaling, have been proposed as pathogenic mechanisms (5, 11, 12), the precise molecular basis of *RELA*-associated immune dysregulation remains incompletely understood.

We investigated six patients from five unrelated families presenting with overlapping autoinflammatory and autoimmune features, all harboring truncating *RELA* mutations. Our findings demonstrate that these mutations lead to a loss of NF-κB signaling function, disrupting the finely tuned balance between NF-κB-mediated pro-survival/pro-inflammatory and cell death pathways. This imbalance results in a shift toward apoptosis and necroptosis, ultimately driving hyperinflammatory responses.

## Results

### Autoinflammation and autoimmunity in patients with truncating RELA mutations

We studied six patients from five unrelated families (**Figure 1**) who presented with signs of autoinflammation and autoimmunity of varying degrees of severity (**Table 1**). P1 presented at the age of 3 years with recurrent fevers, arthralgias, and painful erythematous nodules on the head, the face and the back. Histology of a nodule biopsy was consistent with panniculitis. At the age of 5, she developed lichenoid ulcerating skin lesions in the genital area and osteitis of the head, neck, and mandibular joints. Laboratory tests showed elevated levels of erythrocyte sedimentation rate (ESR), C-reactive protein (CRP), interleukin 6 (IL-6), soluble IL-2 receptor (sIL-2R), serum amyloid a (SAA) and calprotectin. She also had hypergammaglobulinemia, elevated antinuclear antibodies (ANA; 1:160 to 1:1100) and an elevated interferon (IFN) score. Immunophenotyping showed an expansion of IgM^++^CD38^++^ transitional B cells and normal T cells. Genetic testing revealed a heterozygous nonsense variant in the *RELA* gene (NM_021975.4; c.592C>T, p.Arg198*) (**Figure 1A**). The variant was also identified in the patient’s father (P2), who developed autoimmunity at the age of 26 years with vitiligo on the legs, trunk, and face. Like his daughter, he also had an elevated IFN signature. P3 presented with inflammatory symptoms at the age of 15 years, with recurrent oral and genital ulcers, malar rash, arthralgia of the wrists, interphalangeal joints, knees and back, and intermittent abdominal pain with diarrhea and elevated calprotectin. A lesional skin biopsy showed inflammatory infiltrates, vasculitis and microabscesses compatible with Behçet’s disease. Laboratory findings included intermittent lymphopenia with unremarkable lymphocyte differentiation, hypergammaglobulinemia, elevated ANA (1:1280 to 1:20480) and SS-B/La antibodies (strongly positive), and an increased IFN score. Genetic testing revealed a heterozygous *de novo RELA* variant (c.1515_1516del; p.Ala507Profs*25) resulting in a frameshift and premature stop codon (**Figure 1A**). P4 developed recurrent fevers from the age of 18 months associated with fatigue, intermittent abdominal pain and diarrhea, and oral aphthae. Laboratory tests repeatedly showed increased CRP, sIL-2R, SAA, calprotectin, and ANA (1:160). Immunophenotyping showed increased γδ T cells, CD38^++^CD138^+^ plasma cells, and CD19^+^CD38^++^CD138^-^ plasma blasts, indicative of autoimmunity. Exome analysis of P4 revealed a heterozygous *de novo RELA* variant (c.1114C>T, p.Gln372*). P5 presented with severe protein-losing enteropathy at 3 months of age. He also developed refractory atopic eczema from the age of 5 months, which progressed to generalized erythroderma. Laboratory investigations revealed an iron deficiency anemia, hypalbuminemia, hypogammaglobulinemia with normal B cell counts, and IgE-mediated sensitization to several foods. He also had hypothyroidism requiring hormone replacement. Treatment with dupilumab (IL-4/IL-13 antibody) led to a significant improvement of the skin, which still required intensive topical treatment with pimecrolimus and prednicarbate. On a hypoallergenic amino acid-based formula, enteropathy and weight gain improved. Genetic testing identified a heterozygous *RELA* nonsense variant (c.506C>G, p.Ser169*). P6 developed fever episodes and skin rashes in the genital area at 10 months of age, along with elevated CRP. Whole exome sequencing revealed a heterozygous *de novo RELA* variant (c.592C>T, p.Arg198*), which had also been detected in the unrelated patients P1 and P2. None of the patients showed an increased susceptibility to infections.

**Figure 1.**
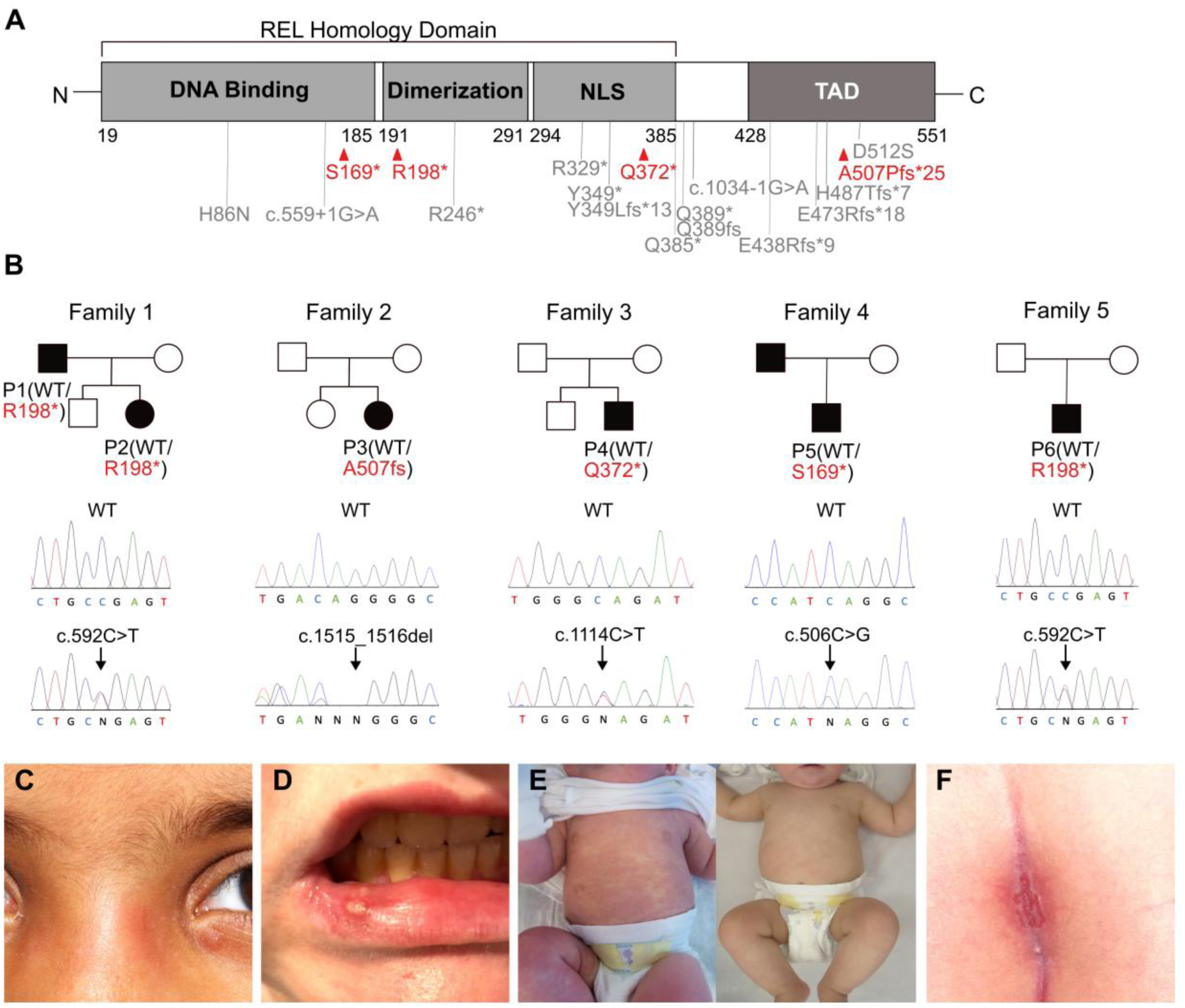
Genetic and clinical findings in patients with truncating *RELA* variants. **(A)** Schematic p65 domain architecture. NLS, nuclear localization signal; TAD, transactivation domain. Newly identified (red) and previously described (grey) pathogenic *RELA* variants. **(B)** Pedigrees and sequence pherograms depicting *RELA* variants. **(C)** Erythema nodosum in periorbital region of P2. **(D)** Oral aphthous ulcer in P3. **(E)** Generalized refractory eczema in P5 and marked clinical improvement under systemic dupilumab. There was no clinical data available for the father of P5. **(F)** Perianal ulcer in P6.

**Table 1.**
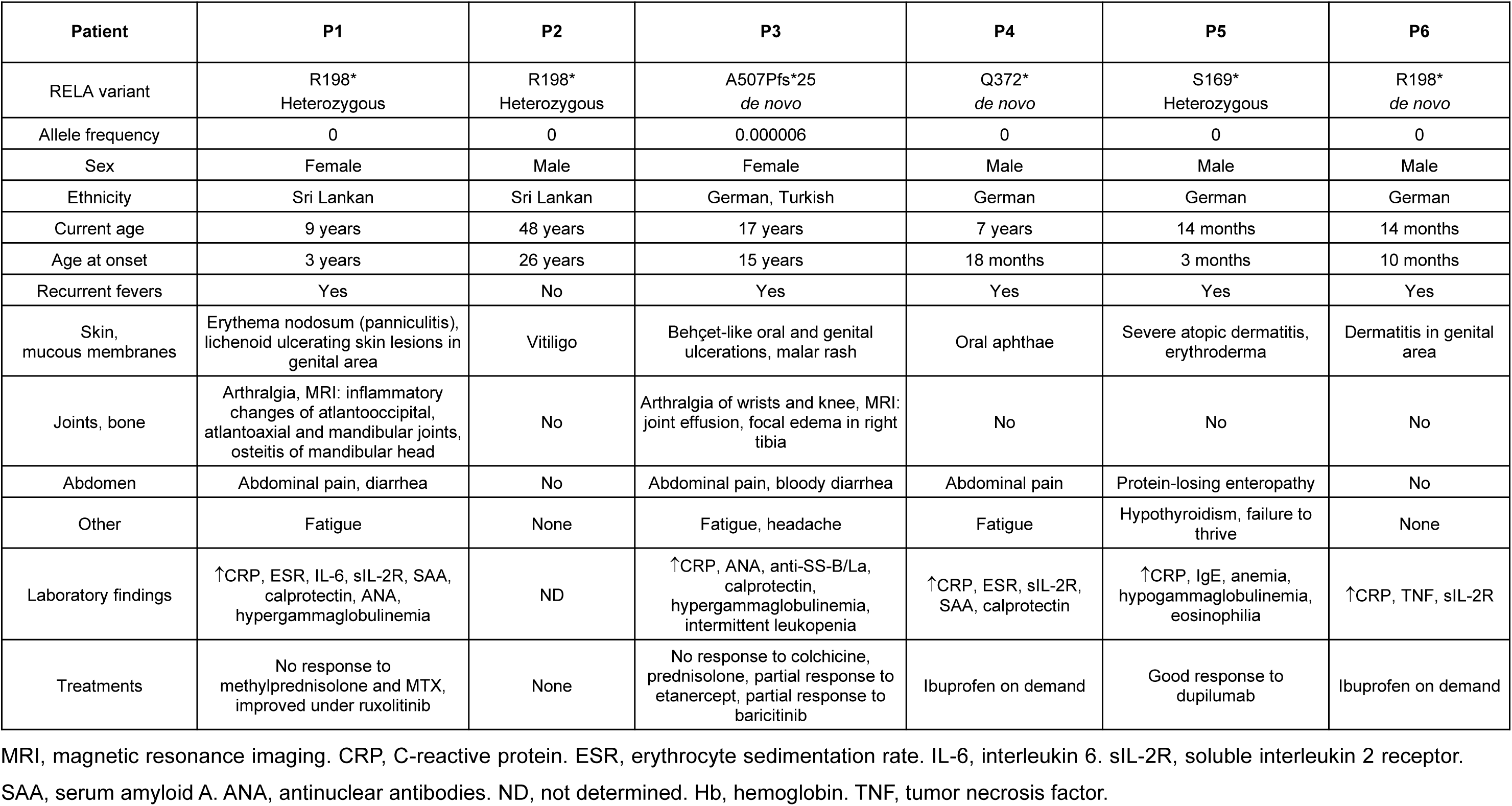
Clinical findings in patients with truncating *RELA* mutations.

| Patient | P1 | P2 | P3 | P4 | P5 | P6 |
| --- | --- | --- | --- | --- | --- | --- |
| RELA variant | R198*<br>Heterozygous | R198*<br>Heterozygous | A507Pfs*25<br>*de novo* | Q372*<br>*de novo* | S169*<br>Heterozygous | R198*<br>*de novo* |
| Allele frequency | 0 | 0 | 0.000006 | 0 | 0 | 0 |
| Sex | Female | Male | Female | Male | Male | Male |
| Ethnicity | Sri Lankan | Sri Lankan | German, Turkish | German | German | German |
| Current age | 9 years | 48 years | 17 years | 7 years | 14 months | 14 months |
| Age at onset | 3 years | 26 years | 15 years | 18 months | 3 months | 10 months |
| Recurrent fevers | Yes | No | Yes | Yes | Yes | Yes |
| Skin, mucous membranes | Erythema nodosum (panniculitis), lichenoid ulcerating skin lesions in genital area | Vitiligo | Behçet-like oral and genital ulcerations, malar rash | Oral aphthae | Severe atopic dermatitis, erythroderma | Dermatitis in genital area |
| Joints, bone | Arthralgia, MRI: inflammatory changes of atlantooccipital, atlantoaxial and mandibular joints, osteitis of mandibular head | No | Arthralgia of wrists and knee, MRI: joint effusion, focal edema in right tibia | No | No | No |
| Abdomen | Abdominal pain, diarrhea | No | Abdominal pain, bloody diarrhea | Abdominal pain | Protein-losing enteropathy | No |
| Other | Fatigue | None | Fatigue, headache | Fatigue | Hypothyroidism, failure to thrive | None |
| Laboratory findings | ↑CRP, ESR, IL-6, sIL-2R, SAA, calprotectin, ANA, hypergammaglobulinemia | ND | ↑CRP, ANA, anti-SS-B/La, calprotectin, hypergammaglobulinemia, intermittent leukopenia | ↑CRP, ESR, sIL-2R, SAA, calprotectin | ↑CRP, IgE, anemia, hypogammaglobulinemia, eosinophilia | ↑CRP, TNF, sIL-2R |
| Treatments | No response to methylprednisolone and MTX, improved under ruxolitinib | None | No response to colchicine, prednisolone, partial response to etanercept, partial response to baricitinib | Ibuprofen on demand | Good response to dupilumab | Ibuprofen on demand |

MRI, magnetic resonance imaging. CRP, C-reactive protein. ESR, erythrocyte sedimentation rate. IL-6, interleukin 6. sIL-2R, soluble interleukin 2 receptor. SAA, serum amyloid A. ANA, antinuclear antibodies. ND, not determined. Hb, hemoglobin. TNF, tumor necrosis factor.

All identified *RELA* variants had not previously been described in the context of an inflammatory disease phenotype and were absent from the gnomAD v4.1.0 database, except for A507Pfs*25 with a very low allele frequency of 0.000006 (**Table 1**). Given a loss of function intolerance (pLI) score of 1 for the *RELA* gene (https://gnomad.broadinstitute.org/gene/-ENSG00000173039?dataset=gnomad_r4), all identified variants were predicted to be deleterious. Consistent with previous reports (13), fibroblasts from P1 and P2 carrying the R198* *RELA* mutation showed expression of mutant p65, while wild type p65 expression was reduced compared to cells from healthy controls (**Supplemental Figure 1A**). As expected, the p65^R198*^ mutant, which lacks both the RHD domain required for dimerization and the transactivation domain, was unable to form homodimers with wild type p65 or heterodimers with p50 and could not transactivate an NF-κB reporter gene (**Supplemental Figure 1, B and C**).

### Increased levels of pro-inflammatory cytokines at base line and in response to immune stimulation

Consistent with clinical signs of autoinflammation and autoimmunity, cytokine profiling of patient sera revealed elevated levels of a broad range of pro-inflammatory cytokines including IL-6, IL-8 IL-10, TNF, MCP-1 (CCL2), and IFN-γ compared to healthy controls (**Figure 2, A-E**). Notably, the inflammasome-dependent cytokines IL-1β and IL-18 (P1-P3) and the Th2-dependent cytokine IL-33 (P2), were also markedly elevated at baseline (**Figure 2, F-H**). In addition, all analyzed patients showed evidence of constitutive type I IFN activation, as shown increased serum levels of IFN-λ (P1-P4), elevation of the type I IFN-dependent chemokine CXCL10 (P1, P2, P5) (**Figure 2, I and J**), or an increased blood IFN signature (P1-P3) (**Figure 2K**), in line with previous reports (11).

**Figure 2.**
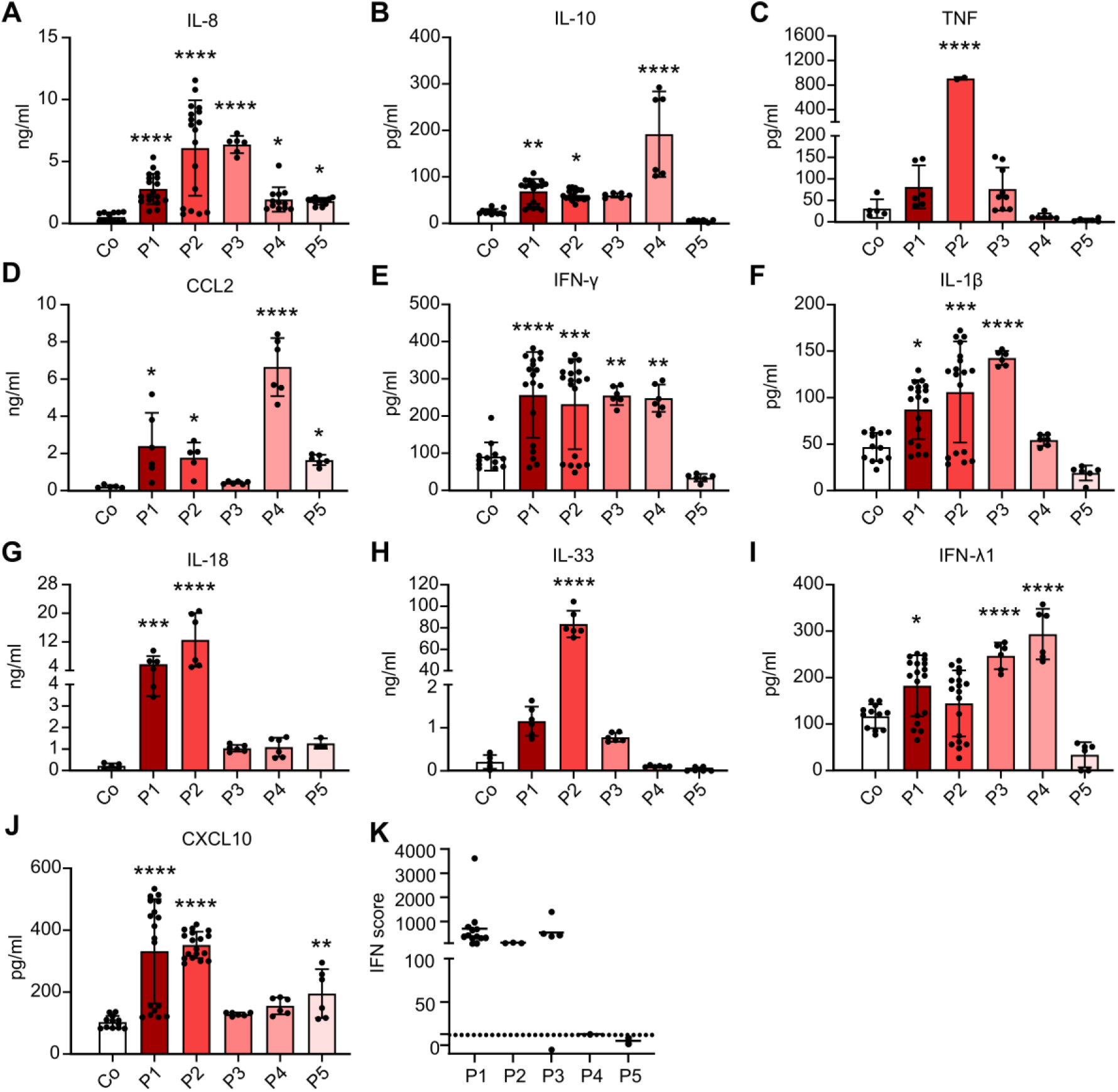
Increased secretion of pro-inflammatory cytokines in *RELA*-perturbed patients. (A-J) Serum levels of IL-8, IL-10, TNF, CCL2, IFN-γ, IL-1β, IL-18, IL-33, IFN-λ1, and CXCL10, collected at different time points, from patients (P1-P5) compared to four healthy controls (Co). Data are plotted as mean ± SD, \**P* < 0.05; \*\**P* < 0.01; \*\*\**P* < 0.001; \*\*\*\**P* < 0.0001 versus mean of wild type controls, Kruskal-Wallis test (Dunn’s multiple comparison test) for IL-8, CCL2 and IL-18 and one-way ANOVA (Dunnett’s multiple comparison test) for IL-10, TNF, IFN-γ, IL-1β, IL-33, IFN-λ1, and CXCL10. **(K)** IFN scores based on expression of IFN-stimulated genes in PBMCs. An IFN score of 12.49 (dashed line) indicates the median IFN score of 10 healthy controls + 2 SD.

To further assess induced cytokine responses, we conducted whole blood stimulation assays using different immunostimulatory ligands. Following stimulation with increasing doses of the TLR4 agonist LPS, all examined patients (P1-P5) secreted significantly higher amounts of IL-6 already at the lowest dose of 1 ng/ml, consistent with hypersensitivity to LPS, whereas at higher doses of 2.5 and 5 ng/ml, patients P3 to P5 showed increased IL-6 secretion (**Figure 3A**). Similarly, TNF secretion was significantly increased by LPS stimulation in a dose-dependent manner in the patients (**Figure 3B**). Notably, all patients also secreted higher amounts of IL-1β in response to LPS compared to controls, indicating hypersensitivity to inflammasome activation (**Figure 3C**). In addition, all patients showed increased IL-10 secretion upon LPS stimulation (**Figure 3D**). Similar to blood cells, primary fibroblasts from patient P1 were hyperresponsive to TNF stimulation compared to a wild type control, as evidenced by a greater induction of the *IL6* gene (**Supplemental Figure 2A**). We next examined the patientś response to stimulation of the type I IFN-inducing nucleic acid sensors TLR7 and TLR9, which have recently been shown to be hyperreactive in patients with truncating *RELA* mutations (11). Upon stimulation of whole blood with increasing doses of the TLR7 agonist R837, two (P4, P5) out of five patients responded with increased secretion of CXCL10, whereas CXCL10 secretion from patients P1, P2, and P3 did not differ from wild type controls (**Figure 3E**). Similarly, after stimulation with the TLR9 agonist ODN2006, only P4 showed an increased CXCL10 response compared to wild type controls (**Figure 3F**). Thus, in our cohort, constitutive type I IFN activation observed in all patients was not consistently associated with TLR7 and TLR9 hypersensitivity. Collectively, these findings demonstrate a global activation of multiple inflammatory pathways in patients with truncating *RELA* mutations that is accompanied by hyperresponsiveness to stimulation with LPS and TNF, and, to a lesser extent, to stimulation of TLR7 and TLR9.

**Figure 3.**
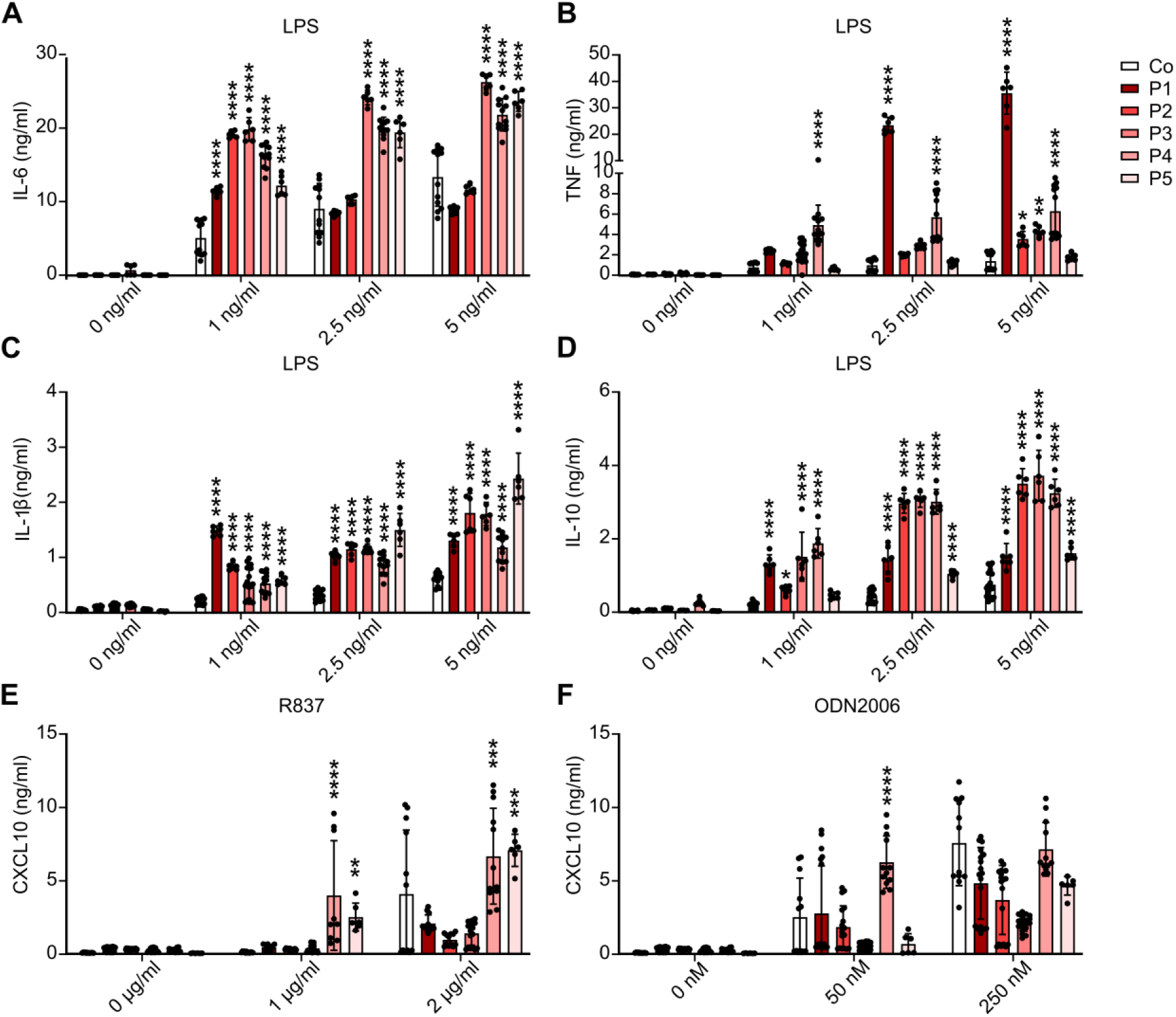
Enhanced cytokine responses following stimulation. (A-D) Secretion of IL-6, TNF, IL-1β, and IL-10 by whole blood from patients (P1-P5) after 24 h stimulation with increasing concentrations of the TLR4 agonist LPS, compared to four healthy controls (Co). Means ± SD. \**P* < 0.05; \*\**P* < 0.01; \*\*\**P* < 0.001; \*\*\*\**P* < 0.0001 versus mean of wild type controls, two-way ANOVA (Dunnett’s multiple comparison test). **(E-F)** Secretion of CXCL10 by whole blood from patients after 24 h stimulation with increasing doses of the TLR7 agonist R837 or the TLR9 agonist ODN2006, compared to four healthy controls. Data are plotted as mean ± SD. \**P* < 0.05; \*\**P* < 0.01; \*\*\**P* < 0.001; \*\*\*\**P* < 0.0001 versus mean of wild type controls, two-way ANOVA (Dunnett’s multiple comparison test).

### Altered canonical NF-κB signaling in patient cells

The finding of upregulated pro-inflammatory cytokines in patient blood, both at steady state and in response to stimuli utilizing the canonical NF-κB pathway, is surprising given the inability of mutant p65 to perform its core function of dimer formation and gene transcription activation (**Supplemental Figure 1, B and C**). To investigate this further, we first examined the subcellular levels of p65 in patient cells. To distinguish wild type from mutant p65, we used an antibody that recognizes the C-terminus and therefore only wild type p65. Confocal imaging revealed reduced levels of cytosolic and nuclear wild type p65 in P1 compared to the control (**Figure 4, A and B**), both in unstimulated cells and after TNF stimulation, consistent with monoallelic expression of wild type p65. A ∼50% decrease of wild type p65 in mutant cells was confirmed by Western blot analysis of cytosolic and nuclear fractions (**Figure 4, C and D**).

**Figure 4.**
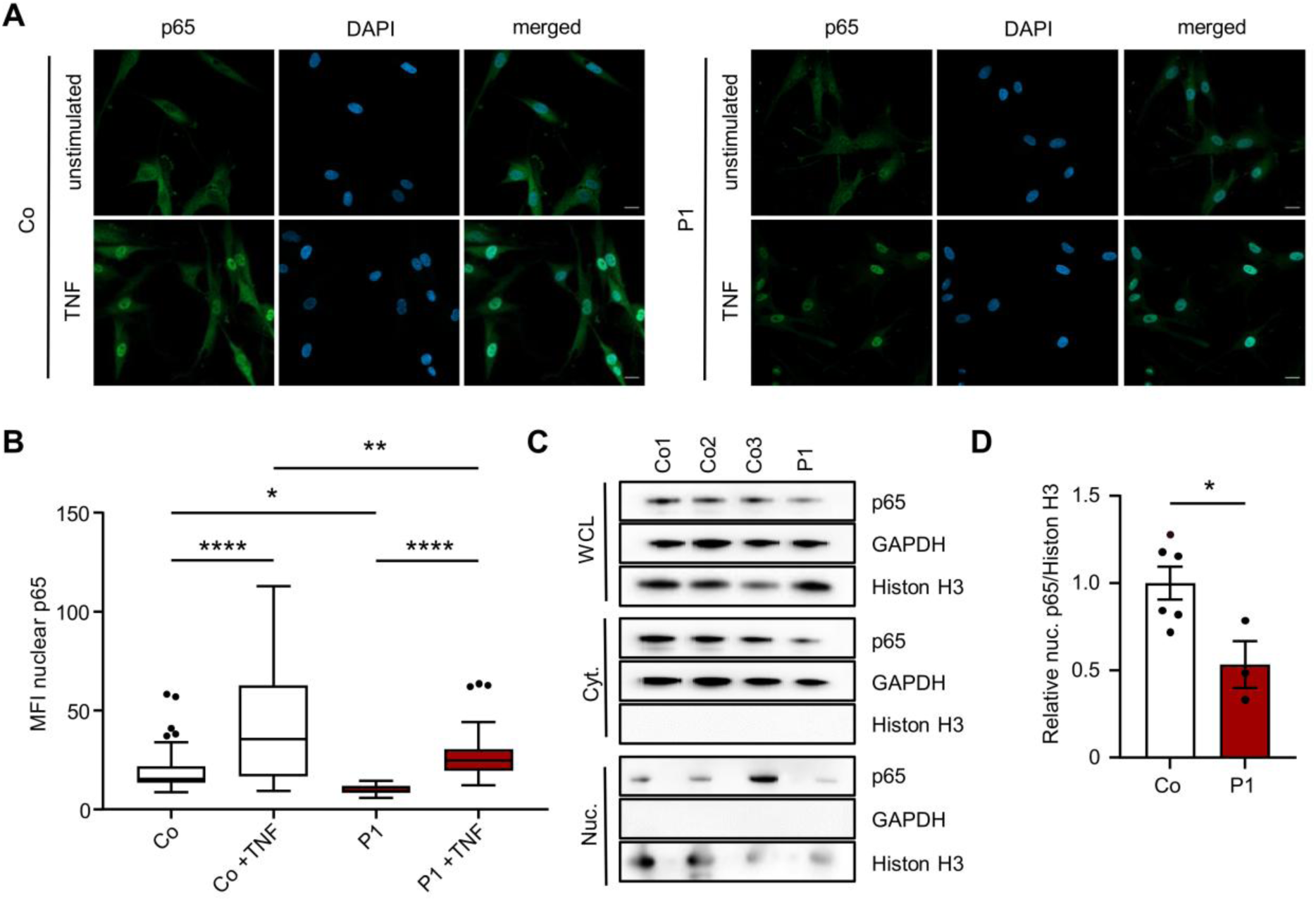
Altered nuclear translocation of p65 in patient fibroblasts. **(A)** Representative images of fibroblasts from patient and control immunostained for p65 (green) without and after TNF-stimulation (10 ng/ml, 1h). Nuclei were stained with DAPI. Scale bars: 20 μm. **(B)** Quantitative analysis of nuclear p65 intensity. Mean fluorescence intensity (MFI) in patient cells (n=30-35) versus wild type control cells (n=28-32). \**P* < 0.05; \*\**P* < 0.01; \*\*\**P* < 0.001; \*\*\*\**P* < 0.0001 versus mean of wild type controls. One-way ANOVA (Šidák’s multiple comparison test) was used for statistical analyses. Data presented as Tukey box plots. **(C)** Representative immunoblot of p65 expression in whole cell lysate (WCL), cytosolic (Cyt.) and nuclear fractions (Nuc.) of fibroblasts from P1 and three wild type controls (Co1-3) after subcellular fractionation. GAPDH (cytoplasmic) and Histon H3 (nuclear) were used as loading controls. **(D)** Quantification of relative nuclear p65 in patient fibroblasts (P1) compared to wild type controls, normalized to histone H3. Means ± SEM of at least three independent experiments. \**P* < 0.05 versus controls, Mann-Whitney test.

We next analyzed canonical NF-κB signaling in lymphocytes from patients P1 and P2 by flow cytometry. We first examined the degradation of the inhibitory IκBα protein as the initial step in the signaling cascade leading to the release of the activated NF-κB dimer into the nucleus. Remarkably, in the unstimulated situation, IκBα protein, which is tightly regulated by the canonical NF-κB pathway, was lower in naïve B, CD4^+^, and CD8^+^ T cells from both patients compared to healthy controls (**Figure 5A**). After stimulation of naïve B cells with anti-IgM and phorbol 12-myristate 13-acetate (PMA) and of naïve CD4^+^ and CD8^+^ T cells with PMA, respectively, the degradation of IκBα measured at 20 minutes post-stimulation did not differ between patients and healthy controls (**Figure 5A and Supplemental Figure 3A**), suggesting normal stimulus-induced initiation of canonical NF-κB signaling. We then measured the activation-induced phosphorylation of p65. Consistent with the observed lower levels of p65 in patient cells at basal state (**Figure 4, A and B**), the levels of phosphorylated p65 (pSer529) were also lower compared to healthy controls in both naïve B cells stimulated with anti-IgM or PMA and naïve CD4^+^ and CD8^+^ T cells stimulated with PMA (**Figure 5B and Supplemental Figure 3B**). Thus, while the basal expression of p65 is reduced, its phosphorylation in response to stimulation occurs normally.

**Figure 5.**
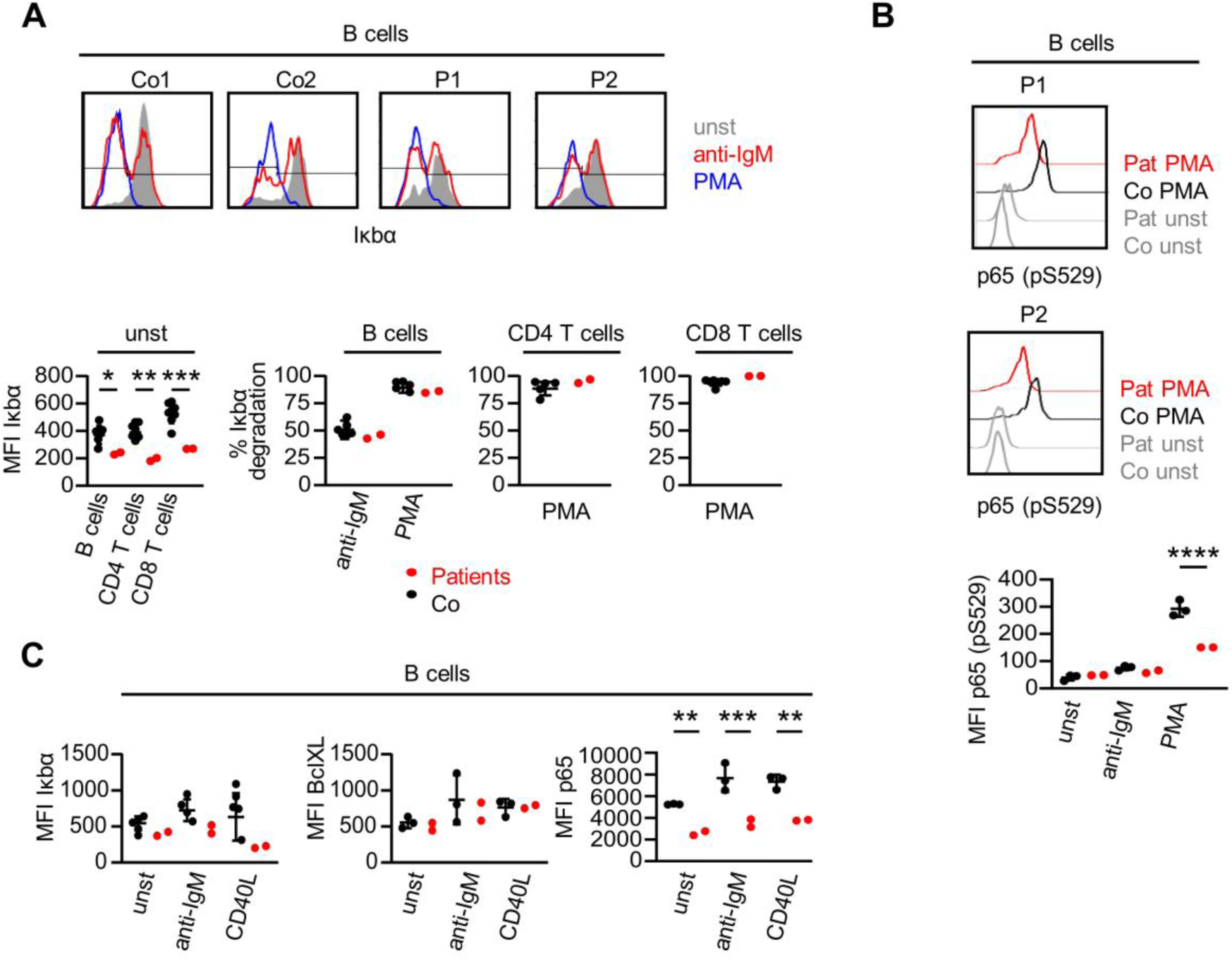
Altered NF-kB signaling in patient lymphocytes. **(A)** IκBα degradation in primary naïve B cells stimulated with anti-IgM and PMA, and in T cells stimulated with PMA, of P1 and P2. IκBα protein levels at baseline and post-stimulation in naïve B, CD4^+^, and CD8^+^ T cells compared to healthy controls (Co). One-way ANOVA (Šidák’s multiple comparison test); \**P* < 0.05; \*\**P* < 0.01; \*\*\**P* < 0.001 versus mean of wild type controls. **(B)** Mean fluorescence intensity (MFI) of phosphorylated p65 upon activation with PMA in naïve B cells of P1 and P2 compared to healthy controls. One-way ANOVA (Šidák’s multiple comparison test); \*\*\*\**P* < 0.0001 versus mean of wild type controls. **(C)** Protein levels of p65 and its targets IκBα (*NFKBIA)* and BclXL (*BCL2L1)* after stimulation of PBMCs of P1 and P2 with anti-IgM and CD40L for 36 h. One-way ANOVA (Šidák’s multiple comparison test); \*\**P* < 0.01; \*\*\**P* < 0.001 versus mean of wild type controls.

To assess the effects of canonical NF-κB pathway stimulation on downstream targets, we stimulated PBMCs with anti-IgM and CD40L and then measured the protein levels of IκBα (NFKBIA) and Bcl-XL (BCL2L1) in naïve B cells after 36 hours. Notably, while IκBα protein levels were adequately induced in B cells from healthy controls, they remained low in patient cells (**Figure 5C and Supplemental Figure 3C**). In contrast, post-stimulation induction of Bcl-XL, which acts as anti-apoptotic protein, in patient cells was not different from that in controls (**Figure 5C and Supplemental Figure 3C**). Consistent with the monoallelic expression of functional p65, p65 protein levels in both unstimulated and stimulated cells were greatly reduced in patient B cells (**Figure 5C and Supplemental Figure 3C**). Taken together, these results indicate that activation of canonical NF-κB signaling upstream of p65 occurs normally in patient cells, albeit at lower levels. However, p65 activation is not accompanied by adequate post-stimulation induction of IκBα which could impair NF-κB inhibition at steady state and result in imbalanced NF-κB signaling.

### Impaired negative feedback control of NF-kB in patient cells and *RELA* gene-edited HEK293T cells

The transcriptional activity of NF-κB is tightly repressed by IκB proteins, including IκBα, IκBβ, and IκBε, through the formation of stable IκB-NF-κB complexes (2). Following stimulus-induced IκB degradation, one of the earliest NF-κB transcriptional targets to be re-expressed is IκBα, providing negative feedback control (14). Consistent with this IκBα-dependent negative feedback mechanism, transfection of wild type p65 into HEK293T cells resulted in a strong induction of IκBα (**Figure 6A**). In contrast, transfection of mutant p65^R198*^ had no effect on IκBα expression (**Figure 6A**), in agreement with its inability to transactivate a reporter gene (**Supplemental Figure 1C**).

**Figure 6.**
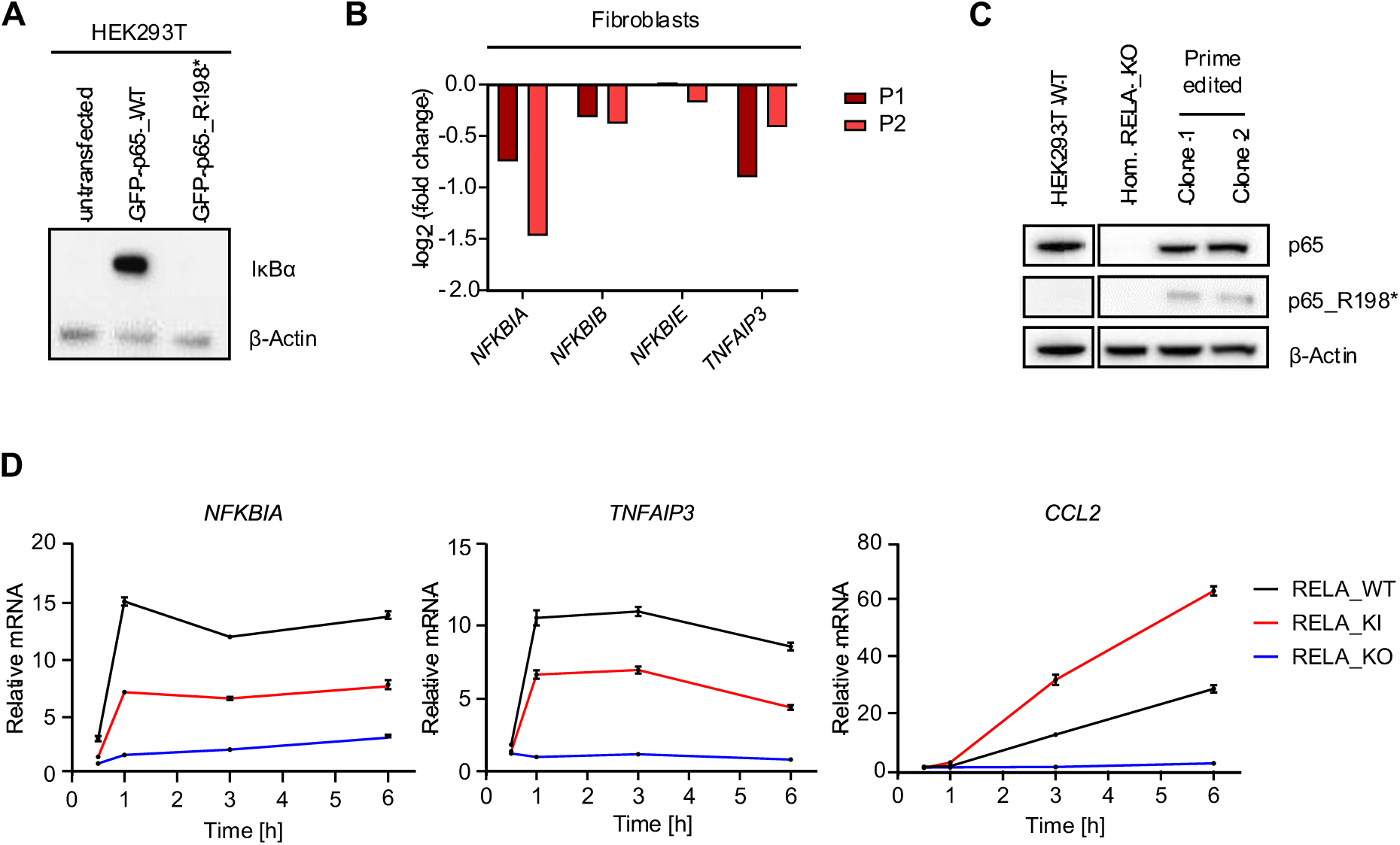
Impaired negative feedback control of NF-kB in patient cells and *RELA*^R198*^ knock-in cells. **(A)** IκBα protein expression in HEK293T cells transfected with either GFP-tagged wild type p65 (GFP-p65_WT) or mutant p65^R198*^ (GFP-p65_R198*). β-actin was probed as loading control. **(B)** Differential gene expression of NF-κB inhibitors *NFKBIA* (IκBα), *NFKBIB* (IκBβ), *NFKBIE* (IκBε), and *TNFAIP3* (A20) in patient fibroblasts (P1, P2), shown as log_2_ fold change compared to three wild type controls. **(C)** Representative immunoblot of p65 expression in wild type HEK293T cells (HEK293T WT) compared to homozygous RELA knockout (RELA_KO) and two independent heterozygous RELA^R198*^ knock-in (RELA_KI) clones (Clone1 and 2). **(D)** Relative expression of *NFKBIA, TNFAIP3,* and *CCL2* in wild type HEK293T cells (RELA_WT), RELA knockout (RELA_KO) and RELA^R198*^ knock-in cells (RELA_KI) after stimulation with 100 ng/ml TNF for indicated time periods. Representative data from one out of three independent experiments run in tricplicates. Data are plotted as mean ± SEM.

Notably, at steady state, IκBα levels also strongly influence basal NF-κB activity by controlling the levels of nuclear NF-κB, which actively shuttles between the nucleus and cytosol. While free IκBα is intrinsically unstable and rapidly degraded in an IKK-independent manner, NF-κB-bound IκBα is more stable (15). Thus, besides regulating its transcription, NF-κB also determines the fate of its own inhibitor IκBα through stabilization. Given the inability of mutant p65 to induce IκBα in HEK293T cells (**Figure 6A**), we reasoned that reduced steady-state levels of p65 might affect basal transcription of IκB protein-coding genes. We therefore examined the consequences of monoallelic expression of functional p65 on the expression of the *NFKBIA* (IκBα), *NFKBIB* (IκBβ), and *NFKBIE* (IκBε) genes in fibroblasts from patients P1 and P2. RNA sequencing revealed a significant decrease in the expression of all three NF-κB inhibitors in the patient fibroblasts compared to three wild type controls (**Figure 6B**), mirroring the reduced basal levels of IκBα observed in unstimulated patient lymphocytes (**Figure 5A**). Moreover, gene expression of the NF-κB inhibitor *TNFAIP3* was also reduced in patient fibroblasts (**Figure 6B**). Thus, at steady state, loss of a functional *RELA* allele in patient cells causes a shift in the balance between NF-κB activating and inhibitory factors toward activation.

To further investigate the functional consequences of the reduction of NF-κB inhibitory factors at the cellular level, we turned to a reductionist approach. We generated HEK293T cells (RELA_WT) with either a complete homozygous knockout of the *RELA* gene (RELA_KO) or with a heterozygous knock-in of the R198* mutation at the *RELA* locus (RELA_KI) by CRISPR/Cas or prime-editing, respectively (**Figure 6C**). Introduction of the R198* *RELA* mutation into HEK293T cells had no effect on cytokine gene expression in unstimulated cells (**Supplemental Figure 4A**). After stimulation of cells with 100 ng/ml of TNF, expression of the NF-κB inhibitory gene *NFKBIA* was strongly induced within 60 minutes in a *RELA* gene dose-dependent manner. Thus, the highest transcriptional activation of *NFKBIA* was seen in RELA_WT cells and the lowest in RELA_KO cells, while RELA_KI cells showed an intermediate transcriptional response (**Figure 6D**), confirming the observation made in patient lymphocytes (**Figure 5A**) that a lack of *RELA* gene dose results in a correspondingly reduced induction of its inhibitor IκBα. Consistent with complete absence of functional p65 in RELA_KO cells, TNF stimulation had essentially no effect on the expression of *TNFAIP3* and *CCL2*, two *bona fide* NF-kB target genes (**Figure 6D**). Normally, TNF-induced expression of *TNFAIP3*, which functions as a key negative regulator of NF-κB signaling, peaks within 1 hour with a kinetic similar to *NFKBIA* (16). However, in contrast to RELA_WT cells, the induction of *TNFAIP3* was significantly reduced in RELA_KI cells, reflecting a weaker transactivation potential exerted by a single *RELA* allele (**Figure 6D**). Interestingly, in RELA_KI cells, in which the induction of the inhibitory genes, *NFKBIA* and *TNFAIP3*, was only half that of RELA_WT cells, the pro-inflammatory *CCL2* gene was strongly induced by more than 2.5-fold at 3 hours and more than 2.2-fold at 6 hours compared to RELA_WT cells (**Figure 6D**). Collectively, these findings suggest that the heightened pro-inflammatory response of RELA_KI cells to TNF stimulation may be due to a lack of inhibitory control of the NF-κB pathway.

### Up-regulation of cell type-specific pro-inflammatory and cell death pathways in patient cells

To further investigate the transcriptional changes related to truncating *RELA* mutations in a cell type-resolved, untargeted manner, we performed single-cell RNA sequencing (scRNAseq) in PBMCs from patients P1, P2, and P3 along with five sex- and age-matched healthy controls. We identified and annotated all major cell types (naïve and memory B cells, naïve and central memory CD4^+^ and CD8^+^ T cells, γδ T cells, NK cells, classical and non-classical monocytes, DCs and pDCs) using annotation transfer and marker-based curation (**Figure 7A and Supplemental Figure 5A**). In line with clinical and laboratory data (**Figure 2K**), cells from all three patients expressed IFN-stimulated genes more highly than control cells - a signal mainly driven by myeloid cells but also present to a lesser degree in lymphoid cells (**Supplemental Figure 5B**). Cell type composition varied strongly with no consistent differences between patient and control samples (**Supplemental Figure 5C**).

**Figure 7.**
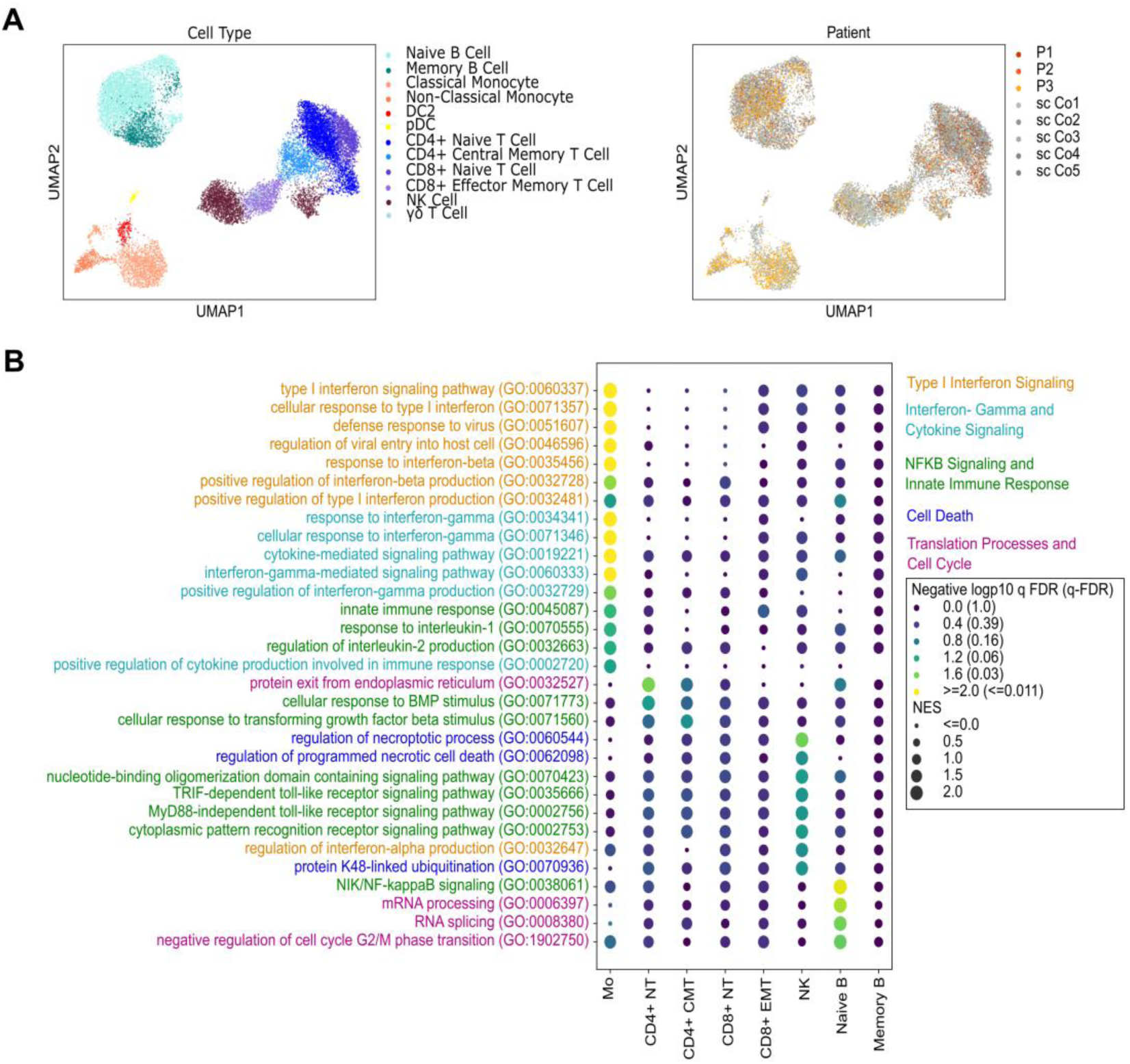
Cell type-specific transcriptomic alterations in *RELA*-perturbed patient cells. **(A)** UMAP embedding of all cells profiled with scRNAseq from patient (P1-P3) and control (scCo1-scCo5) PBMCs. **(B)** Gene Set Enrichment Analysis (GSEA) enrichment of selected Gene Ontology (GO) Biological Processes terms in differentially expressed genes between patient and control cells of the indicated cell types. FDR, False Discovery Rate; NES: Normalized Enrichment Score.

Consistent with the hyperinflammatory phenotype observed in patients, gene set enrichment analysis revealed strong activation of multiple pathways related to pro-inflammatory cytokine production and signaling, including type I IFN, IFN-γ, and IL-1, particularly in monocytes (**Figure 7B**). Among the top-ranked gene sets enriched in monocytes were “type I interferon signaling pathway” (GO:0060337) and “response to interferon-gamma” (GO:0034341) (**Figure 7B and Figure 8A**). The IFN-stimulated genes, *ISG15* and *APOBEC3G*, as well as the antiviral factor *BST2*, which can amplify pro-inflammatory responses via NF-κB signaling, showed increased and relatively homogeneous expression in both classical and non-classical monocytes (**Figure 8B and Supplemental Figure 5D**). In contrast, the pro-inflammatory cytokine *CCL2* and *NFKBIA*, both NF-kB target genes, were more prominently upregulated in classical monocytes (**Figure 8B**), potentially indicating more wild type-like NF-kB activity in classical monocytes.

**Figure 8.**
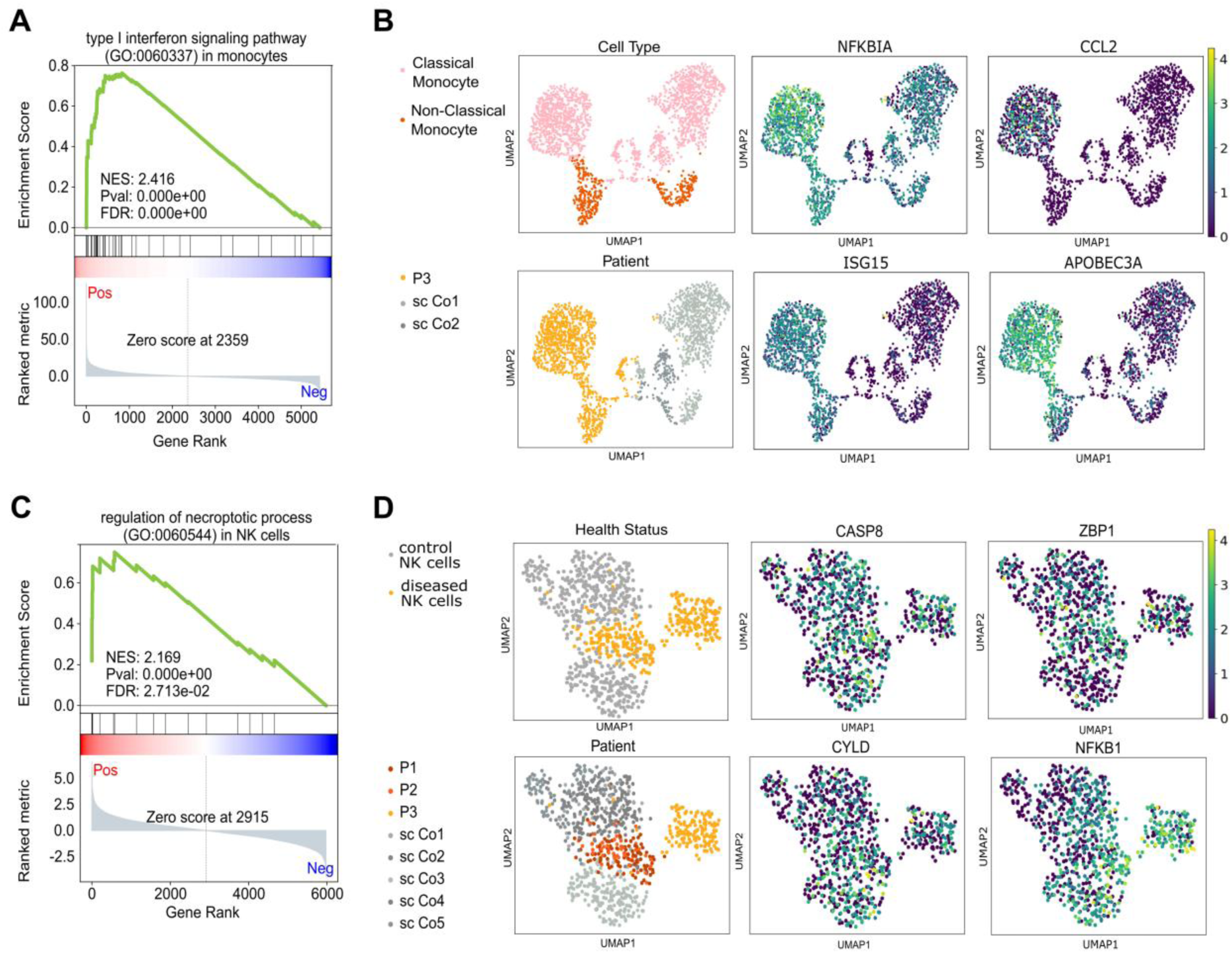
Activation of inflammatory and cell death pathways in *RELA*-perturbed patient cells. **(A)** Gene Set Enrichment Analysis (GSEA) leading edge plot for the GO term “type I interferon signaling pathway” enriched in monocytes from patient (P1-P3) and control (scCo1-scCo5) PBMCs. **(B)** UMAP embedding of monocytes colored by the expression of the indicated genes in individual cells. **(C)** Gene Set Enrichment Analysis (GSEA) leading edge plot for the GO term “regulation of necroptotic process” enriched in NK cells. **(D)** UMAP embedding of NK cells colored by the expression of the indicated genes in individual cells.

In naïve B cells from patients, “NIK/NF-κB signaling”(GO:0038061) emerged as the most significantly enriched pathway (**Figure 7B**), suggesting a negative regulatory crosstalk of p65 on the alternative NF-kB signaling pathway in circulating B cells. Transcription factor activity analysis, considering the expression of all known target genes for a given transcription factor, further revealed elevated activity scores for *STAT1*, *NFKB*, and *RELA* in patient B cells (**Supplemental Figure 6A**). Patient B cells also showed elevated expression of the NF-κB inhibitors and target genes (*NFKBIA*, *NFKBID*, and *NFKBIE*), unlike patient T cells (**Supplemental Figure 6B**), providing evidence of constitutive NF-κB pathway activation in B cells.

In contrast, patient T and NK cells, in which IκBα protein levels were more strongly reduced than in B cells (**Figure 5A**), showed upregulation of cell death pathways, including both apoptosis and necroptosis (**Figure 7B, 8C, and Supplemental Figure 6C**), the latter being a lytic, pro-inflammatory form of programmed cell death. This signature was most pronounced in NK cells (**Figure 7B and 8C**). “Apoptosis” as enriched term was also evident in naïve CD4⁺ and CD8⁺ T cells, but not in monocytes or B cells (**Supplemental Figure 6C**). Interestingly, genes driving the apoptosis signal in T cells and NK cells were slightly different, yet highly different to those driving the signal in monocytes (**Supplemental Figure 6D**). Notably, *CASP8* and *ZBP1* emerged among the most upregulated differentially expressed genes in NK cells (**Figure 8D and Supplemental Figure 6E**). *CASP8* encodes caspase-8, a critical initiator of apoptosis downstream of death receptor signaling, while *ZBP1* (Z-DNA binding protein 1) functions as a cytosolic nucleic acid sensor that can activate both necroptosis and apoptosis. Through these mechanisms, ZBP1 contributes to inflammation via the release of danger-associated molecular patterns (DAMPs) (17, 18). Accordingly, the highest-ranked gene set enriched in patient NK cells was “regulation of necroptotic process” (GO:0060544) (Figure 7B). This enrichment was amongst others driven by increased expression of *CASP8*, *ZBP1,* and *CYLD*, and *JUN* was also among the top differentially expressed genes (**Figure 8D and Supplemental Figure 5D and 6E**), consistent with chronic activation of both apoptotic and necroptotic cell-death pathways. In addition, transcription factor activity analysis revealed the highest activity scores for *JUN*, *ATF4*, and *CREB1* in CD4^+^ naïve T cells from patients (**Supplemental Figure 6A**), reflecting a state of activation and stress-associated dysfunction. Jun is associated with T cell activation, promotes Th1 and Th17 differentiation, and has been linked to T cell exhaustion (19). ATF4 acts as a central mediator of the integrated stress response and regulates the expression of pro-inflammatory cytokines under cellular stress conditions (20). CREB1 contributes to the transcription of key cytokines such as IL-2 (essential for T cell proliferation), IFN-γ (critical for Th1 responses), and under certain conditions, IL-17 (involved in Th17 differentiation) (21). Together, these transcriptional signatures support a bias toward pro-inflammatory effector T cells, primarily Th1 and Th17, which are known to play key roles in initiating and sustaining chronic inflammation. Overall, the cell type-resolved transcriptional comparison of patient versus control immune cells uncovered a differential signature marked by reduced NF-κB inhibition and heightened apoptotic/necroptotic signaling, particularly in T and NK cells.

### Enhanced sensitivity to TNF-induced apoptosis and necroptosis in patient cells and *RELA* gene-edited HEK293T cells

Upon TNF receptor engagement, adaptor proteins including RIPK1 and cIAP1/2 (cellular inhibitor of apoptosis proteins) assemble complex I, a membrane-associated platform where ubiquitinated RIPK1 activates NF-κB and promotes survival (22). Deubiquitination of RIPK1 by CYLD, A20, or OTULIN shifts signaling toward cytosolic complex II, composed of FADD, procaspase-8, and RIPK1, where caspase-8 activation initiates apoptosis and simultaneously suppresses necroptosis by cleaving RIPK1 and RIPK3. In the absence or inhibition of caspase-8, RIPK1 and RIPK3 interact via RHIM domains to form the necrosome, which drives inflammatory necroptosis, particularly in response to TNF (23).

Given the increased apoptosis/necroptosis signaling (**Figure 7B and Supplemental Figure 6C**), upregulation of *CASP8* and *CYLD* (**Figure 8D and Supplemental Figure 6, D and E**), and a reduction in negative regulation of NF-κB signaling (**Figure 5A**), observed predominantly in patient T and NK cells, we hypothesized that patient cells may be intrinsically more sensitive to cell death. To test this, we stimulated patient cells with TNF in the presence of the SMAC mimetic birinapant, which inhibits cIAPs and promotes apoptosis (24). In a parallel condition, cells were stimulated with TNF and co-treated with birinapant and the pan-caspase inhibitor emricasan, to block apoptosis and enforce necroptosis (25). Upon treatment of fibroblasts from patients P1 and P2 with birinapant, TNF stimulation led to a stronger induction of cleaved caspase-3, indicating enhanced apoptosis compared to wild type controls (**Figure 9A**). Similarly, co-treatment with birinapant and emricasan resulted in markedly increased phosphorylation of RIPK1 at Ser166, a hallmark of necroptosis activation, compared to wild type controls, suggesting heightened necroptotic potential in patient cells (**Figure 9A**). Notably, RELA_KI cells, expressing only one functional *RELA* allele, also exhibited increased apoptosis and necroptosis in response to TNF stimulation compared to RELA_WT or RELA_KO cells (**Figure 9B**), confirming that *RELA* haploinsufficiency underlies the heightened sensitivity to TNF-induced cell death observed in patient cells. Collectively, these findings demonstrate that the loss of a functional *RELA* allele compromises TNF tolerance and predisposes cells to inflammatory cell death.

**Figure 9.**
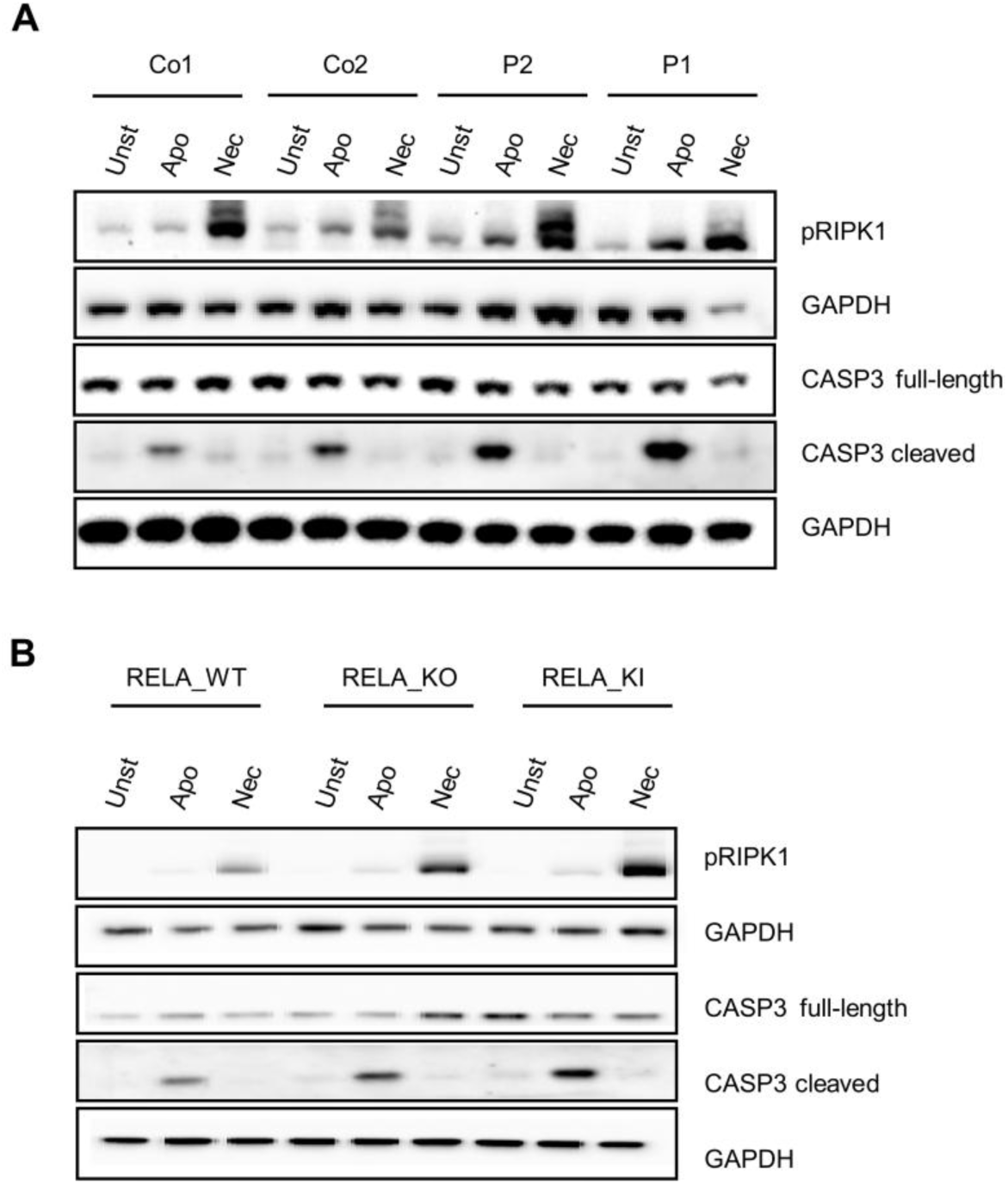
Increased sensitivity to apoptosis and necroptosis conferred by the *RELA*^R198*^ mutation. **(A)** Increased phosphorylated RIPK1 (pRIPK1) and cleaved caspase-3 (CASP3) in patient fibroblasts (P1 and P2) compared to two wild type controls (Co1 and Co2) upon TNF stimulation. Apo: stimulated with TNF and co-treated with birinapant; Nec: stimulated with TNF and co-treated with birinapant and emricasan. GAPDH was stained as loading control. **(B)** Increased phosphorylated RIPK1 (pRIPK1) and cleaved CASP3 in wild type (RELA_WT), *RELA* knockout (RELA_KO), and *RELA*^R198*^ knock-in (RELA_KI) HEK293T cells. GAPDH was probed as loading control.

## Discussion

This study delineates the clinical, immunological, and mechanistic consequences of heterozygous truncating mutations in *RELA*, encoding the NF-κB subunit p65. By combined analysis of patient-derived cells and gene-edited heterologous cell models, we demonstrate that these variants confer *RELA* haploinsufficiency. This state of reduced gene dosage impairs the negative regulatory control of NF-κB signaling, leading to insufficient induction of inhibitory feedback regulators such as IκBα and A20. As a consequence, the tightly regulated interplay between the pro-survival and pro-death functions of NF-κB becomes compromised, shifting immune homeostasis toward immune dysregulation and ultimately driving self-sustaining hyperinflammation.

We identified four novel truncating *RELA* mutations in six patients presenting with a spectrum of autoinflammatory and autoimmune disease of varying severity. All patients exhibited inflammatory skin disease, including Behçet-like mucosal ulcerations, erythema nodosum, erythroderma, and malar rash. Five out of six patients showed systemic autoinflammatory features, such as recurrent fever and elevated inflammatory markers, while four patients developed intestinal inflammation, consistent with previous reports of RELA-associated disease (5–8, 11–13). Autoimmune manifestations such as vitiligo, hypothyroidism, autoimmune anemia, and the presence of autoantibodies further highlight the dual autoinflammatory-autoimmune phenotype associated with truncating *RELA* variants.

All identified mutations were either inherited in an autosomal dominant manner or occurred *de novo* and disrupted the transactivation domain of p65 required for transcriptional activity. Consistent with this, the R198* mutant failed to form homodimers or p65/p50 heterodimers, abolishing its core functional capacity. Remarkably, patient P3, who carried the most C-terminal truncating mutation reported to date (A507Pfs), exhibited a clinical phenotype highly similar in both symptoms and severity to that of patient P1 with the more N-terminal R198* mutation. This indicates that impairment of the transactivation domain alone is sufficient to confer a loss-of-function phenotype.

Mechanistically, *RELA* haploinsufficiency primarily impairs negative feedback regulation rather than upstream NF-κB activation. Despite preserved stimulus-induced phosphorylation of p65, patient cells failed to adequately induce IκBα and other inhibitory regulators such as A20, resulting in impaired termination of NF-κB signaling. *RELA^R198*^*knock-in HEK293T cells recapitulated this phenotype, showing reduced induction of inhibitory regulators but enhanced and prolonged expression of pro-inflammatory cytokines such as CCL2. Importantly, these amplified inflammatory responses depended on the residual activity of the functional *RELA* allele and not on the mutant protein, underscoring a gene-dosage effect in which one functional *RELA* allele seems to be sufficient to activate inflammatory target genes but insufficient to maintain proper negative feedback control of NF-κB signaling.

While prompt activation of NF-κB is essential for host defense, uncontrolled NF-κB signaling can be detrimental and impede immune homeostasis, promoting chronic inflammation. Among its negative regulatory factors, IκB proteins play a central role in limiting the intensity and duration of NF-κB responses (14–16). In our patients, p65 levels were consistently reduced, in line with monoallelic expression, while basal IκBα levels differed across cell types. With otherwise preserved stimulus-induced NF-κB activation, insufficient induction of IκBα and A20 emerged as the critical defect disturbing the negative feedback loop that normally constrains NF-κB activity at steady state.

Single-cell transcriptomics revealed that the consequences of this impairment were cell type-specific: Monocytes showed strong enrichment of type I IFN, IFN-γ, and IL-1 signaling pathways, with increased expression of *ISG15*, *APOBEC3G*, and *BST2*. B cells exhibited enhanced *RELA* activity and alternative NF-κB signaling, which warrants further investigation but may reflect compensatory mechanisms supporting survival and antibody function (26). By contrast, T cells and NK cells, which exhibited markedly reduced IκBα levels, were skewed toward apoptotic and necroptotic programs, characterized by increased expression of *CASP8*, *ZBP1*, and *CYLD*, alongside increased activity of stress-related transcription factors *JUN*, *ATF4*, and *CREB1*. This transcriptional landscape indicates that *RELA* insufficiency renders lymphocytes particularly vulnerable to stress-induced and inflammatory cell death, thereby fueling chronic inflammation through the release of DAMPs (27). Consistently, patient fibroblasts and RELA_KI cells demonstrated enhanced apoptosis and necroptosis upon TNF stimulation, reflecting heightened susceptibility to inflammatory cell death. These findings are consistent with prior work in RELA-deficient mice and fibroblasts, which demonstrated profound TNF sensitivity and tissue injury (5, 28, 29). Notably, while complete p65 deficiency in mice causes embryonic lethality due to extensive hepatocyte apoptosis, heterozygous *RELA* knockout mice remain phenotypically normal (30). The discrepancy between heterozygous *RELA* deficiency in mice and the pronounced phenotype in humans may reflect differences in microbial and inflammatory exposures, which constantly challenge the NF-κB regulatory capacity in humans but not in mice maintained under pathogen-free conditions. The marked clinical variability among patients may similarly reflect differences in environmental triggers that aberrantly activate NF-κB signaling.

In conclusion, we demonstrate that truncating *RELA* mutations impair NF-κB feedback control, resulting in heightened inflammatory signaling, TNF sensitivity, and increased apoptotic and necroptotic cell death. This mechanistic framework explains the diverse clinical features observed in patients with truncating *RELA* mutations and suggests new therapeutic avenues aimed at restoring NF-κB feedback regulation and limiting inflammatory cell death.

## Methods

### Sex as a biological variable

The patients in our study were male and female patients, sex was not considered as a biological variable. However, when comparing patients with healthy controls, samples were matched for sex and age.

### Exome sequencing

Exome sequencing was performed as part of the routine diagnostic work-up for patients with suspected monogenic immune dysregulation. Genomic DNA was isolated from peripheral blood leukocytes, and whole-exome sequencing was carried out in accredited diagnostic laboratories following standard protocols. Sequence data were aligned to the human reference genome (GRCh37/hg19 or GRCh38, depending on the center) and analyzed for rare, protein-altering variants in genes associated with immune-related disorders. Candidate variants were confirmed by Sanger sequencing and assessed for segregation within families when parental samples were available. Identified variants were classified according to the guidelines of the American College of Medical Genetics and Genomics (ACMG).

### Sanger sequencing

Genomic DNA sequences flanking *RELA* (NM_021975.4) gene mutations were amplified by PCR using gene-specific primers (Eurofins MWG Operon) and sequenced in both directions using the BigDye Terminator v1.1 Cycle Sequencing Kit (Thermo Fisher Scientific, 4337449) on a 3130xl Genetic Analyzer (Applied Biosystems). Data were analyzed using the Vector NTI software (Life Technologies).

### Cell culture and stimulation

Primary fibroblasts obtained from patients and healthy controls (passages 4 to 20) and HEK293T cells were cultured in Dulbecco’s modified Eagle’s medium (DMEM; Sigma-Aldrich, D6546) supplemented with 2 mM L-glutamine (Gibco, 25030-024), 1% antibiotics/antimycotics (Gibco, 15240-062), 5% non-essential amino acids (NEAA, Gibco, 11140-035) and 10% fetal bovine serum (FBS, Sigma-Aldrich, S0615) at 37°C under 5% CO_2_. Cells were treated with 10 or 100 ng/ml TNF (Prospec Protein Specialists, 112PTNFA31). For isolation of human PBMCs, whole blood diluted with phosphate-buffered saline (PBS), was gently layered over an equal volume of BioColl (Sigma-Aldrich) and centrifuged for 30 min at 400x g without brake. The intermediate layer containing PBMCs was removed and added to prewarmed medium.

### Mutagenesis

For mutagenesis, the GFP-RELA plasmid (addgene, 23255) was used. The R198* mutation was introduced by site-directed mutagenesis using the QuikChange Lightning Site-Directed Mutagenesis Kit (Agilent Technologies, 210518) and the following primers: fwd: CGAGCTCAAGATCTGCTGAGTGAACCGAAACTC and rev: GAGTTTCGGTTCACTCAGC-AATCTTGAGCTCG (Eurofins Genomics). GFP was replaced by mCherry using AgeI and BSrG1 High Fidelity restriction enzymes.

### Western blot analysis

The cells were collected and washed twice with PBS. Pellets were lysed in radioimmunoprecipitation assay (RIPA) buffer (50 mM TRIS-HCl, pH 7.4, 150 mM NaCl, 1 mM EDTA, 1% Triton X-100, 1 mM sodium orthovanadate, and 20 mM sodium fluoride) supplemented with 1x Complete Protease Inhibitor Cocktail, 1x PhosSTOP phosphatase inhibitors (Roche), and 1 U/ml DNase I (Qiagen, 79254). Protein concentration was determined using a BCA Kit (Thermo Fisher Scientific). Lysates were resolved in a 4 to 12% NuPAGE Bis-TRIS gel (Thermo Fisher Scientific) under reducing and denaturing conditions and blotted onto a nitrocellulose membrane (Sigma-Aldrich). Membranes were blocked in 5% nonfat dry milk (AppliChem, APPA0830,0500) or in 5% BSA (Sigma-Aldrich, 05470) in 1X TBS plus 0.1% Tween-20 (SERVA, 37470.01) and incubated overnight at 4°C using the following antibodies: anti-GAPDH (Cell Signaling Technology, 2118), anti-p65 (Santa Cruz, sc-8008), anti-IkBα (R&D Systems, AF4299), anti-p50 (Thermo Fisher Scientific, MA5-15870), anti-histone H3 (Cell Signaling Technology, 9715S), anti-β-actin (Sigma-Aldrich, A5316) anti-pRIPK1 (Cell Signaling, 65746), anti-CASP3 (Cell Signaling, 9668), and anti-GFP (Cell Signaling, 2555S). Antibodies were diluted in 5% dry milk or 5% BSA in 1X TBS plus 0.1% Tween-20. Immunoreactive signals were detected by chemiluminescence using the SuperSignal West Femto Maximum Sensitivity Substrate (Thermo Fisher Scientific, 34095) on an Azure imaging system. Quantification of imaged bands was performed using ImageJ.

### Co-immunoprecipitation

For co-immunoprecipitation, HEK293T cells were lysed in RIPA buffer supplemented with 1x Complete Protease Inhibitor Cocktail, 1x PhosSTOP phosphatase inhibitors (Roche), and 1 U/ml DNase I. Lysates were incubated with equilibrated G-agarose beads (Thermo Scientific, 53125) for 1.5 h at 4°C and then centrifuged at 2500x g and 4°C for 5 min. The supernatant was removed, and after three washing steps with 500 µL ice-cold PBS and renewed centrifugation, precipitated proteins were eluted and denatured in 80 µL 2x SDS loading buffer, incubated at 95°C for 10 min, and centrifuged (2500x g at 4°C for 2 min). The supernatant removed corresponded to the eluate. SDS-PAGE and Western blotting of the eluted protein samples and signal detection were performed as described above.

### Bulk RNA sequencing

RNA sequencing was performed as previously described (31).

### Single-cell RNA sequencing

Frozen PBMCs from patients 1 (sc P1), 2 (sc P2), and 3 (sc P3), along with 5 age- and sex-matched controls (sc Co1, sc Co2, sc Co3, sc Co4, sc Co5), were rapidly thawed at 37°C and immediately resuspended in 5% FBS/PBS. After centrifugation at 350-550 rcf for 5 min at 4°C, the cell pellets were resuspended and incubated for 10 min with 5 μL of 10% Human TruStain FcX™ Fc Receptor Blocking Solution (BioLegend) in PBS. For sample multiplexing, 48.5 μL of antibody master mix (comprising 47.5 μL of 5% FBS/PBS, 0.5 μL of PE anti-human CD123, and 0.5 μL of APC anti-human CD303 (BDCA-2)) and 1.5 μL of TotalSeq™-C antibody hashtags (BioLegend) were added to each sample. The samples were then incubated at 4°C for 20 min, followed by washing and resuspension in 5% FBS/PBS. Cell counts were assessed using the Invitrogen Countess II Automated Cell Counter, and samples were pooled to ensure equal cell contributions. Pool 1 consisted of patients 1 and 2, and controls sc Co3, sc Co4, and sc Co5, and pool 2 consisted of patient 3 and controls sc Co1 and sc Co2. SYTOX™ Green (Thermo Fisher, S7020) diluted 1:1000 in PBS was added, and dead cells were removed, while pDCs were enriched using the FACS BD FACSAria™ III Cell Sorter (BD Biosciences). To focus on lymphoid cell populations, myeloid cells were filtered out for pool 1, while they were retained for pool2. Following cell sorting, two populations were obtained: (i) pDCs and (ii) remaining PBMCs. Both cell populations were centrifuged, and the pellets were resuspended in 0.1% BSA in PBS - pDCs in 300 µL and the remaining PBMCs in 100 µL. Cell concentration of the PBMC fraction was quantified using the Countess automated cell counter (Thermo Fisher Scientific). The cell concentration was recorded and based on the 10x Genomics recommended loading concentration (33,000 cells for a target recovery of 20,000), the required volume of PBMCs to reach this target was calculated. The 300 µL pDC-only suspension was then divided equally into two 150 µL aliquots. The appropriate volume of sorted PBMCs, calculated to yield a total of 33,000 cells per reaction, was added to each aliquot to generate pDC-only aliquot to generate pDC-enriched cell suspensions. These mixed suspensions were subsequently spun down and the supernatant was carefully removed to leave ≤38.7 µL final volume, consistent with the maximum volume allowable for loading onto the 10x Genomics Chromium platform, and loaded onto two lanes of the Chip G of the Chromium Next GEM Single Cell 5’ v2 (Dual Index) Kit (10x Genomics). GEX libraries were constructed according to the kit instructions. Both libraries were sequenced on a P3 Flow Cell on an Illumina NextSeq 1000 machine.

### Single-cell RNA Data Analysis

Raw sequencing data in FASTQ format were processed to generate a gene-cell expression matrix using the Cell Ranger 7.1.0 count pipeline. The parameters were set with expect-cells = 20,000 and include-introns = True, utilizing the GRCh38-2020-A human genome reference. Cells were demultiplexed using an in-house hashtag demultiplexer and genetically with CellSNP-lite (v1.2.2) (32) and Vireo (v0.5.8) (33). Doublets were identified and subsequently removed using our hashtag demultiplexer, Vireo (v0.5.8) (33) and scDblFinder (v1.8.0) (34). Further analysis was conducted using the Scanpy framework (v1.9.3) (35). Quality filtering was performed using pp.filter_cells(min_genes = 200) and pp.filter_genes(min_cells = 3). Cells with more than 25% of transcripts mapped to the mitochondrial genome were excluded. The filtered dataset consisted of 54,272 cells. Normalization to correct for library size was carried out using the pp.normalize_total(target_sum = 10,000) function, ensuring comparability of counts among cells. The normalized data underwent log-transformation using the pp.log1p function. Highly variable genes were identified using pp.highly_variable_genes and were utilized for dimensionality reduction and clustering. Principal component analysis (PCA) was performed using tl.pca. A neighborhood graph for cells was generated using the pp.neighbors function with n_neighbors = 15. The neighborhood graphs were then embedded using the tl.umap function and visualized through the pl.umap function. Cell type annotation was done using model-based automatic cell type annotation CellTypist (v1.5.0) (36) using the models Immune_all_Low.pkl and Immune_all_High.pkl Annotation was further refined manually using established cell marker genes.

Differential gene expression analysis (DGEA) was done using decoupleR (v.1.6.0) (37) in a pseudobulk manner (except for monocytes). Pseudobulks were generated using dc.get_pseudobulk, which sums up the counts per cell type per patient. Genes were further filtered using dc.plot_filter_by_expr (group=’mutation’, min_count = 10, min_total_count = 15) and the in decoupleR integrated pydeseq2 (v.0.4.9) used for statistical tests. The DESeq2 object was built for each cell type, using DeseqDataSet(refit_cooks = True, inference = DefaultInference) and as design_factors the variable denoting diseased or healthy cells given. To compute log fold changes, dds.deseq2 was run and the contrast extracted with DeseqStats. For hypothesis testing, the Wald test was applied using contrast.summary. Transcription factor activity analysis was also done using decoupleR and their Univariate Linear Model (ULM) decoupler.mt.ulm with CollecTRI (38) as resource for transcriptional regulatory interactions. Because monocytes were only collected from one patient (P1) and the two corresponding controls, a pseudobulk approach as above was not possible, and differential gene expression analysis was done in a single cell manner using scanpy.tl.rank_genes_groups using method = ‘t-test’.

Gene Set Enrichment Analysis (GSEA) was performed using GSEApy (v1.1.4) (39), utilizing gene rankings derived from the differential gene expression analysis based on the t-value. For monocytes, only genes expressed in at least 10% of the diseased cells were considered. For all other cell types, only genes with an average mean expression of at least 10 counts were included. The Human MSigDB Hallmark Gene Sets 2020 (40, 41) and the GO Biological Processes Gene Sets 2021 (42, 43) were tested for enrichment using the function prerank.

### Luciferase-based reporter assay

500,000 cells/well of the NF-kB/293/GFP-LucTM Transcriptional Reporter Cell Line (System Biosciences, TR860A-1) were seeded for transfection the following day. The pEGFP-N1 (GenBank Accession, U55762), GFP-p65_WT, and GFP-p65_R198* plasmids were transfected using the lipofectamineTM 3000 (Invitrogen, L3000-008) reagent following the manufacturer’s manual. Cells were stimulated with 10 ng/µl for 16 hours with TNF (Prospec Protein Specialists, 112PTNFA31). The Luciferase Assay System (Promega, E1500) was used for measurement of luciferase activity according to manufactureŕs instructions. Bioluminescence was quantified using a Mithras LB 940 multi plate reader (Berthold Technologies).

### Immunofluorescence

Cells were fixed with 4% formaldehyde, permeabilized with 0.25% Triton X-100 in PBS for 5 min at room temperature and blocked with 1% BSA in PBS for 1 h at room temperature before overnight incubation with the primary antibody (anti-p65; Invitrogen, 51-0500) diluted in 1% BSA-containing PBS at 4°C. After three washes with PBS, cells were incubated with the appropriate Alexa Fluor-labeled secondary antibody (Invitrogen, A-11008) diluted in 1% BSA-containing PBS for 1 h in the dark at room temperature. Finally, cells were washed for an additional three times with PBS before mounting with VectaShield-containing DAPI. Confocal microscopy was performed using an inverted LSM980 (Zeiss) equipped with an Airyscan detector unit. Images were acquired using a 20×/0.8 plan-apochromat objective with z-stack series at 0.31 μm intervals for 7 sections using the piezo drive, followed by raw image processing using the Airyscan processing and extended depth of focus functions in Zen Black software (Zeiss). Mean fluorescence intensities were determined in individual cells using a custom-written MATLAB code (44).

### Apoptosis and necroptosis assay

Primary human fibroblasts and HEK293T cells were cultured to confluence in DMEM complete medium. For apoptosis and necroptosis induction, TNF (Biolegend, 570108), birinapant (Biomol, Cay19699-5), and emricasan (Biomol, Cay22204-5) were dissolved in DMSO and a master mix was prepared using DMEM complete medium. Cells were exposed to 1 µM birinapant for 5 hours, followed by 20 ng/ml TNF for a further 14 hours to induce apoptosis. To induce necroptosis, cells were first incubated with 1 µM birinapant and 5 µM emricasan for 5 hours, followed by 20 ng/ml TNF for a further 14 hours. Post-stimulation, cells were trypsinized, centrifuged (800 rpm, 5 min), and washed twice with PBS. For downstream analysis, pellets were stored in -80°C. DMSO was used as solvent control.

### Cell fractionation

Frozen cell pellets from cultured fibroblasts were resuspended in ice-cold 0.1% NP40 in PBS. An aliquot of the lysate was removed as “whole cell lysate”, mixed with 6x Laemmli sample buffer and then kept on ice until sonication. The remaining lysate was centrifuged and the supernatant removed as “cytosolic fraction”, mixed with 4x Laemmli sample buffer, and boiled at 95°C for 1 min. The remaining pellet was resuspended in 1 mL of ice-cold 0.1% NP40 in PBS and centrifuged. After discarding the supernatant, the pellet was resuspended with 1x Laemmli sample buffer as “nuclear fraction”. Whole cell lysate and nuclear fraction were sonicated for 5 sec at high level and then boiled for 1 min at 95°C. Cell fractions were verified by Western blot analysis using anti-GAPDH and anti-histone H3 as cytoplasmic and nuclear fraction-specific antibodies, respectively, as described (45).

### Quantitative RT-PCR

RNA was extracted with the ReliaPrep RNA Cell Miniprep System (Promega, Z6012) followed by DNase I digestion. RNA was reverse transcribed using the GoScript Reverse Transcription System (Promega, A5001). Target gene expression was determined by quantitative RT-PCR using GoTaq qPCR Master mix (Promega, A6002) on a QuantStudio 5 Real-Time PCR System (Applied Biosystems). Relative expression of mRNAs was analyzed by the cycle of threshold (Ct) value of target genes and quantified by normalizing to *GAPDH* (fwd: GAAGGTGAAGGTCGGAGTC; rev: GAAGATGGTGATGGGATTTC) and hypoxanthine phosphoribosyltransferase 1 (fwd: AGATGGTCAAGGTCGCAAG; rev: TTCATTATAGTCAAGGGCATATCC) using the ΔΔCt method. The following primers were used: *CXCL10* (fwd: GTGGCATTCAAGGAGTACCTC; rev: TGATGGCCTTCGATTCTGGATT); *MX1* (fwd: GTTTCCGAAGTGGACATCGCA; rev: CTGCACAGGTTGTTCTCAGC); *IL-6* (fwd: AGACAGCCACTCACCTCTTCAG; rev: TTCTGCCAGTGCCTCTTTGCTG); *TNFAIP3* (fwd: CTCAACTGGTGTCGAGA AGTCC; rev: TTCCTTGAGCGTGCTGA ACAGC), *NFKBIA* (fwd: TCCACTCCATCCTGAAGGCTAC; rev: CAAGGACACCAAAAGCTCCACG); *CCL2* (fwd: AGAATCACCAGCAGCAAGTGTCC; rev: TCCTGAACCCACTTCTGCTTGG); *TNF* (fwd: ATGAGCACTGAAAGCATGATCC; rev: GAGGGCTGATTAGAGAGAGGTC).

### Blood IFN score

Total RNA was extracted from PBMCs using the ReliaPrep RNA Cell Miniprep System (Promega, Z6012), followed by DNase I digestion. RNA was reverse-transcribed using the GoScript Reverse Transcription System (Promega, A5001). Gene expression was determined by quantitative RT-PCR using the TaqMan Universal PCR Master Mix (Applied Biosystems, #4427788) on an ABI7300 and normalized to *GAPDH* (fwd: GAAGGTGAAGGTCGGAGTC; rev: GAAGATGGTGATGGGATTTC) and hypoxanthine phosphoribosyltransferase 1 (Hs02800695_m1, Thermo Fisher Scientific) expression. For calibration, a calibrator cDNA was included in each assay. Target genes were analyzed using predesigned TaqMan probes (Thermo Fisher Scientific) for *IFI27* (Hs01086373_g1), *IFI44* (Hs00951349_m1), *IFI44L* (Hs00915292_m1), *IFIT1* (Hs01675197_m1), *ISG15* (Hs01921425_s1), *RSAD2* (Hs01057264_m1), and *SIGLEC1* (Hs00988063_m1). The IFN score was calculated as previously described (46).

### Cytokine analysis

Cytokines were quantified using the LEGENDplex Human Inflammation Panel 1 (BioLegend) and the LEGENDplex Human Anti-Virus Response Panel (BioLegend) according to the manufacturer’s instructions. Data were collected on a LSR Fortessa flow cytometer (BD Biosciences) and analyzed with the LEGENDplex Data Analysis V8.1 software (BioLegend).

### Whole Blood assay

Heparinized blood was distributed in 140 µL aliquots into a 96-well plate and maintained in RPMI medium (Gibco, 31870-025) at 37 °C. For analysis of induced cytokine responses, blood was stimulated with LPS (1, 2.5, or 5 ng/ml; Invivogen, tlrl-b5lps), R837 (1 or 2 µg/ml; Invivogen, tlrl-imqs-1), or ODN2006 (50 or 250 nM; Invivogen, tlrl-2006) for 24 hours at 37°C. After incubation, plates were centrifuged at 800x g for 5 minutes at room temperature. Supernatants were pipetted into a new 96-well plate and frozen at -80°C.

### Intracellular flow cytometry

IκBα degradation and phosphorylation of p65 were determined as described previously with minor adaptations (47, 48). In brief, 3.5 x 10^5^ PBMCs were left untreated or stimulated with 15 µg/ml F(ab)’2 goat anti-human IgM (Southern Biotech) for 35 min, recombinant CD40L, or 50 ng/ml PMA (Sigma Aldrich) for 15 min. Cells were fixed by addition of Cytofix and permeabilized by using Perm III (both BD Biosciences) following the manufactureŕs instructions. After permeabilization, cells were stained with the appropriate antibodies to discriminate T and B cell subsets and measured on a LSR Fortessa (BD Biosciences). For measurement of IκBα, Bcl-xL, and p65, 3 x 10^5^ PBMCs were stimulated with 10 µg/ml anti-human IgM and CD40L for 36 hours. Zombie NIR (Biolegend) was added 10 min prior to fixation with Cytofix and permeabilization with Perm III as described above. After washing, cells were stained with the respective antibodies and measured on a LSR Fortessa. Data were analysed with Flowjo 10.10. The following antibodies were used: anti-CD19 Brilliant Violett 421, anti-CD21 PE-Cy7, anti-CD38 PerCp-Cy5.5, anti-CD45RA BV605, anti-CD3 BV650, anti-CD4 BV786 (BioLegend); anti-CD27 Brilliant Violett 605, anti-CD27 BUV395, anti-CD8 BUV495, anti-p65(pS256) Alexa Fluor 488, anti-IκBα PE, (BD Biosciences); anti-Bcl-xL Alexa Fluor® 488, rabbit anti-human p65 XP AF647 (Cell Signaling Technology); anti-IgD FITC (Southern Biotech); anti-IgM Alexa Fluor 647 (Jackson Immunoresearch Laboratories).

### Generation of RELA_KO and RELA_KI HEK293T cells

For CRISPR/cas9-mediated generation of *RELA* knockout cells, the px459 plasmid pSpCas9(BB)-2A-Puro (addgene, 62988), containing a cutting Cas9, a single-guide (sg) RNA scaffold, and a selection marker, was used for insertion of targeting sequences as described (49). sgRNA targeting the *RELA* gene was constructed using the following oligonucleotides (Eurofins Genomics): fwd: TAATACGACTCACTATAGTGCCGAGTGAACCGAA; rev: TTCTAGCTCT-AAAACGAGTTTCGGTTCACTCGGC. For insertion of the sgRNA into the plasmid, the following guides were used: fwd: CACCGTGCCGAGTGAACCGAAACTC; rev: AAACGAGTTTCGGTTCACTCGGCAC. HEK293T cells were seeded in 6-well plates and transfected with the Px459 plasmid including the inserted sgRNA scaffold using lipofectamine^TM^ 3000 (Thermo Fischer Scientific) in DMEM complete media containing 10 mg/ml puromycin. 24 hours after transfection, medium was replaced with DMEM complete with 1 µg/mL puromycin. After completion of puromycin selection, cells were sorted into a 96-well plate for generation of single clones. For prime editing, a pU6-pegRNA-GG acceptor was used to generate heterozygous *RELA* R198* knock-in cells (RELA_KI). PegRNA templates were synthesized by PCR and then cloned into the pU6-pegRNA-GG-acceptor. Successful editing was verified by Sanger sequencing and by Western blot analysis using anti-p65.

### Statistical analysis

Statistical analysis was carried out in GraphPad Prism 10. For normally distributed variables, parametric tests were used, including *t* test for comparison of two groups and one-way analysis of variance (ANOVA) for comparison of three or more groups. For variables with non-normal distribution, nonparametric tests were used. A *P* value less than 0.05 was considered significant. Data are presented as mean ± SD or mean + SEM as indicated.

## Study approval

The study was conducted in accordance with the Declaration of Helsinki and with approval by the Ethics Committee (EK386102017, BO-EK-466102022) of the Medical Faculty, Technische Universität Dresden. Written informed consent was obtained from the patients or their legal representatives.

## Data availability

RNA sequencing data generated in this study have been deposited in the Single Cell Portal database under accession code SCP2151. All other data are available within the article or the Supplemental Materials.

## Author contributions

NL and MLK conceptualized the research. MLK supervised clinical data curation and experimental research. JK supervised scRNAseq data generation and analysis. NL, SWe, BK, AD, SE, TS, MB, SWa, PS, SK, Skr, and CW conducted the experiments, acquired data, and analyzed results. AE, TR, MM, and AK generated scRNAseq data. SVH, RAJ, and UH conducted exome sequencing and variant analysis. AE performed scRNAseq data analysis with contributions from SM. ÖS, LI, TN, MF, HO, CS, and JKD provided clinical data. JS acquired patient material. MLK wrote the manuscript with contributions from NL, SW, CW, AE, and JK. All authors reviewed and edited the manuscript.

## Acknowledgments

We thank the patients and their families for participation in the study. We acknowledge the assistance of the flow cytometry and the imaging facilities of the Center for Molecular and Cellular Bioengineering and the Medical Theoretical Centre, TU Dresden, and the Lighthouse Core Facility unit of the University Medical Center Freiburg. This work was supported by German Research Foundation (Deutsche Forschungsgemeinschaft) grants LU 2342/1-1 (NL), CRC237 369799452/B21 (MLK, JK), CRC237 369799452/A11 (MLK), CRC369 501752319/C06 (MLK), CRC237 369799452/A06 (CW), and project number 545533246 (BK), and by the German Federal Ministry of Education and Research (Bundesministerium für Forschung, Technologie und Raumfahrt, BMFTR) grant 01GM2206C (GAIN, MLK), 01GM2206A (GAIN, BK) and as part of the German Center for Child and Adolescent Health (DZKJ) under the funding code 01GL2405B (CS, MLK). SW was supported by a DFG Gerok fellowship (CRC237 369799452). JK was supported by an EKFS starting grant (2019_A70). MM was supported by a DAAD PhD fellowship. The LSR Fortessa was funded by the DFG (project number 446167311).

---

**Supplemental Figure 1.**
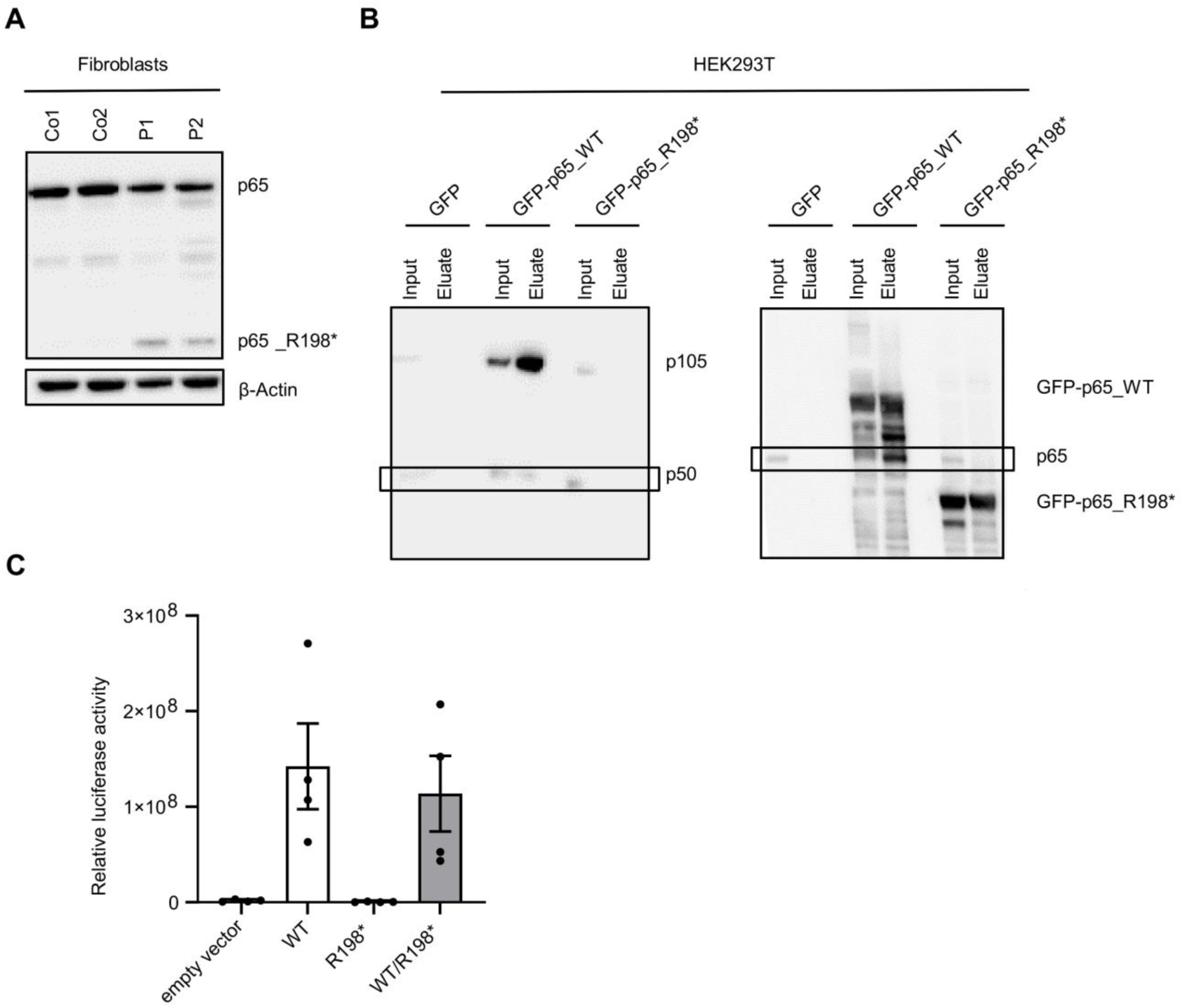
Functional Characterization of the truncating *RELA*^R198*^mutation. **(A)** Expression of wild type (p65) and mutant (p65_R198*) p65 in patient fibroblasts (P1, P2) in comparison to two wild type controls (Co1, Co2). β-Actin was probed as loading control. **(B)** p65 homodimer and heterodimer formation assayed by co-immunoprecipitation of whole-cell lysates from HEK293T cells transfected with either GFP-tagged wild type (GFP-p65_WT) or mutant (GFP-p65_R198*) p65 using anti-p65 antibody, followed by Western blot analysis using anti-p65 (homodimer) or anti-p50 antibody (heterodimer). **(C)** NF-κB activity measured by luciferase reporter assay in HEK293 cells expressing wild type (WT) or mutant (R198*) p65 or both (WT/R198*).

**Supplemental Figure 2.**
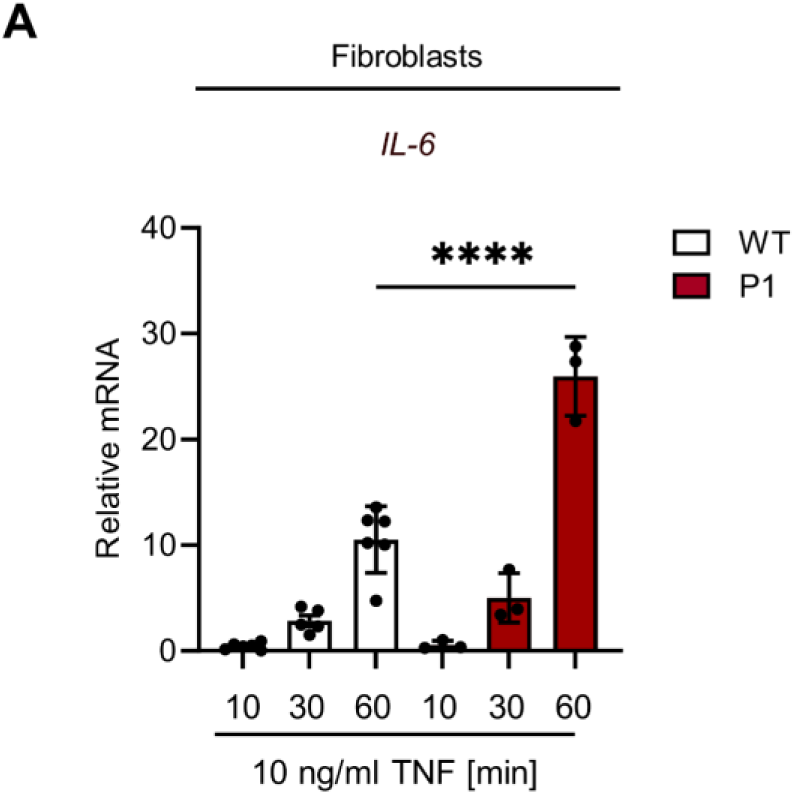
Patient fibroblasts are hyperresponsive to TNF stimulation. **(A)** Relative expression of *IL-6* in primary fibroblasts of patient P1 compared to two healthy controls (WT). Cells were stimulated with 10 ng/ml TNF for the indicated time periods. Data represent mean ± SD of three independent experiments. ****P < 0.0001 versus mean of wild type controls, one-way ANOVA (Šidák’s multiple comparison test).

**Supplemental Figure 3.**
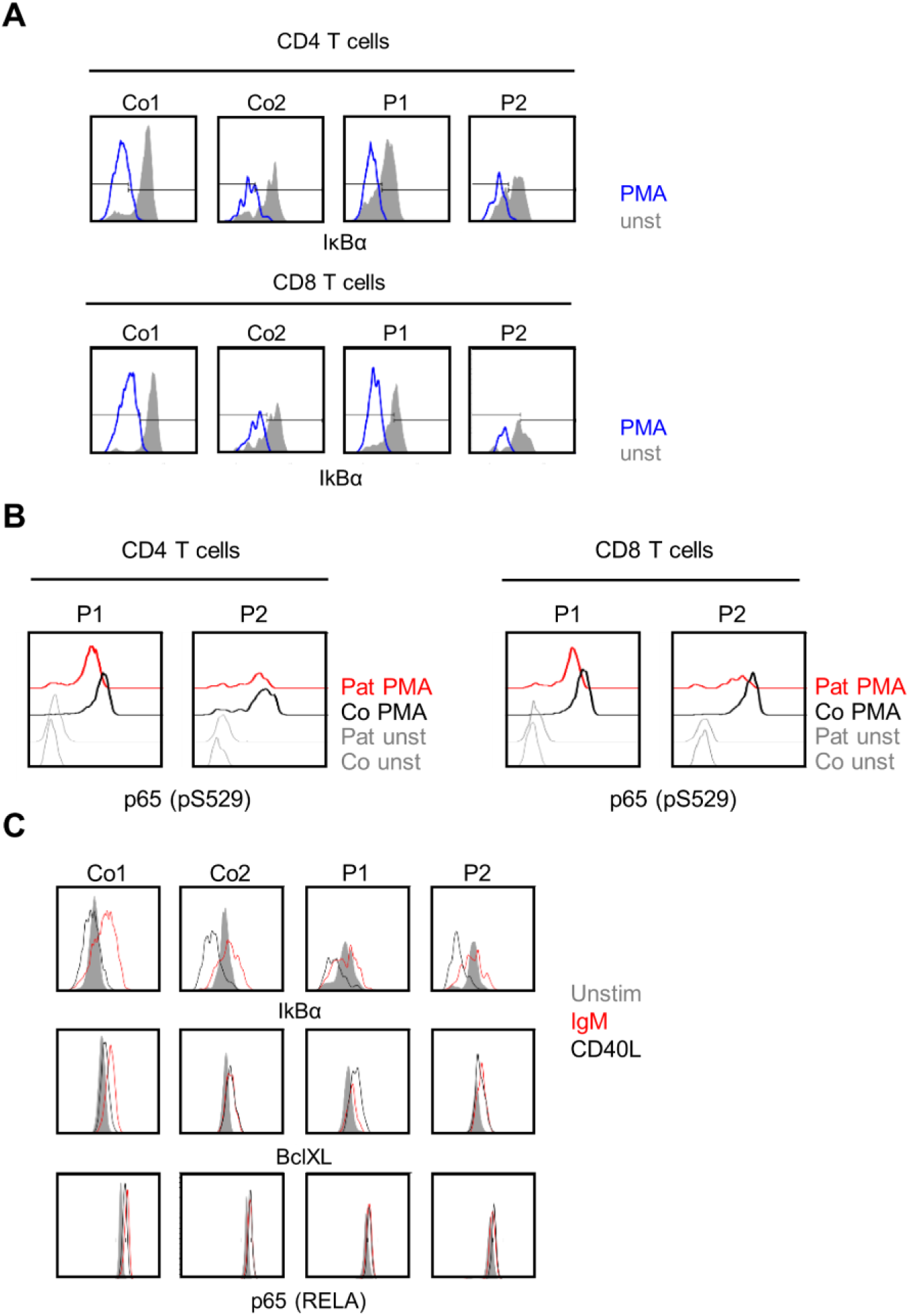
Altered NF-kB signaling in patient PBMCs. **(A)** IκBα degradation in PMA-stimulated CD4^+^ and CD8^+^ T cells from patients (P1, P2) compared to healthy controls (Co1, Co2). **(B)** Reduced p65 phosphorylation (pS529) in patient T cells upon activation of canonical NF-kB signaling. Mean fluorescence intensity (MFI) of phospho-p65 upon activation with PMA in CD4^+^ and CD8^+^ T cells of patients compared to healthy controls. **(C)** Protein expression of p65 and its targets IκBα (*NFKBIA)* and BclXL (*BCL2L1*) in PBMCs of patients and controls after stimulation with anti-IgM and CD40L for 36 h.

**Supplemental Figure 4.**
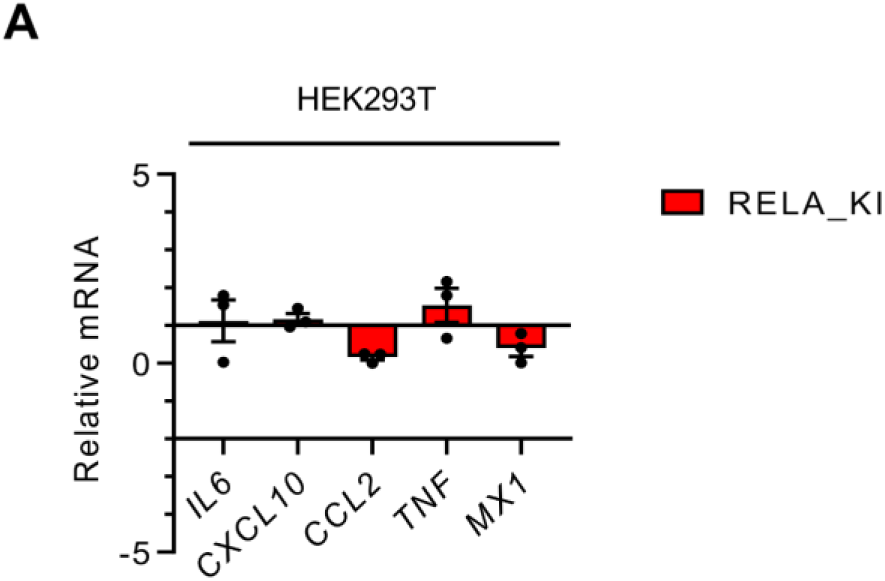
Basal cytokine gene expression in heterozygous *RELA*^R198*^ knock-in HEK293T cells. **(A)** Relative expression of *IL6, CXCL10, CCL2, TNF,* and *MX1* in unstimulated *RELA^R198*^* knock-in HEK293T cells (RELA_KI) in comparison to wild type HEK293T cells, measured by qRT-PCR. Data are plotted as mean ± SD of three independent experiments.

**Supplemental Figure 5:**
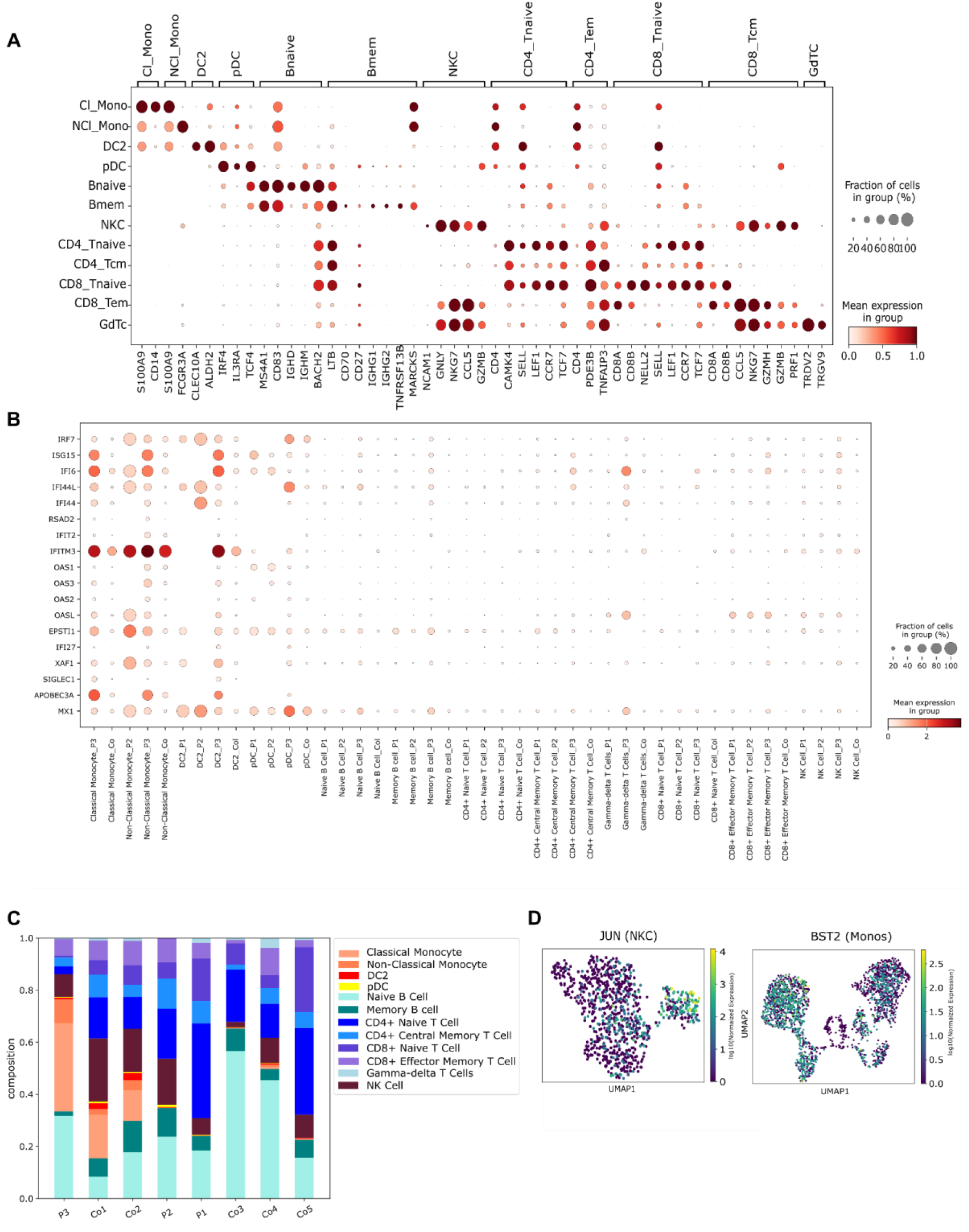
Single-cell RNA seq analysis of PBMCs. **(A)** Dotplot showing the expression of marker genes used to annotate the indicated cell types. **(B)** Dotplot showing the expression of Type I interferon signature genes in patient and control cells of the indicated cell types. **(C)** Barplot showing the cell type composition of patient and control samples. For patient 1 and 2 and controls (sc Co3, sc Co4 and sc Co5) monocytes were removed during FACS. **(D)** UMAP embedding of monocytes and NK cells, respectively, colored by the expression of the indicated genes in individual cells.

**Supplemental Figure 6:**
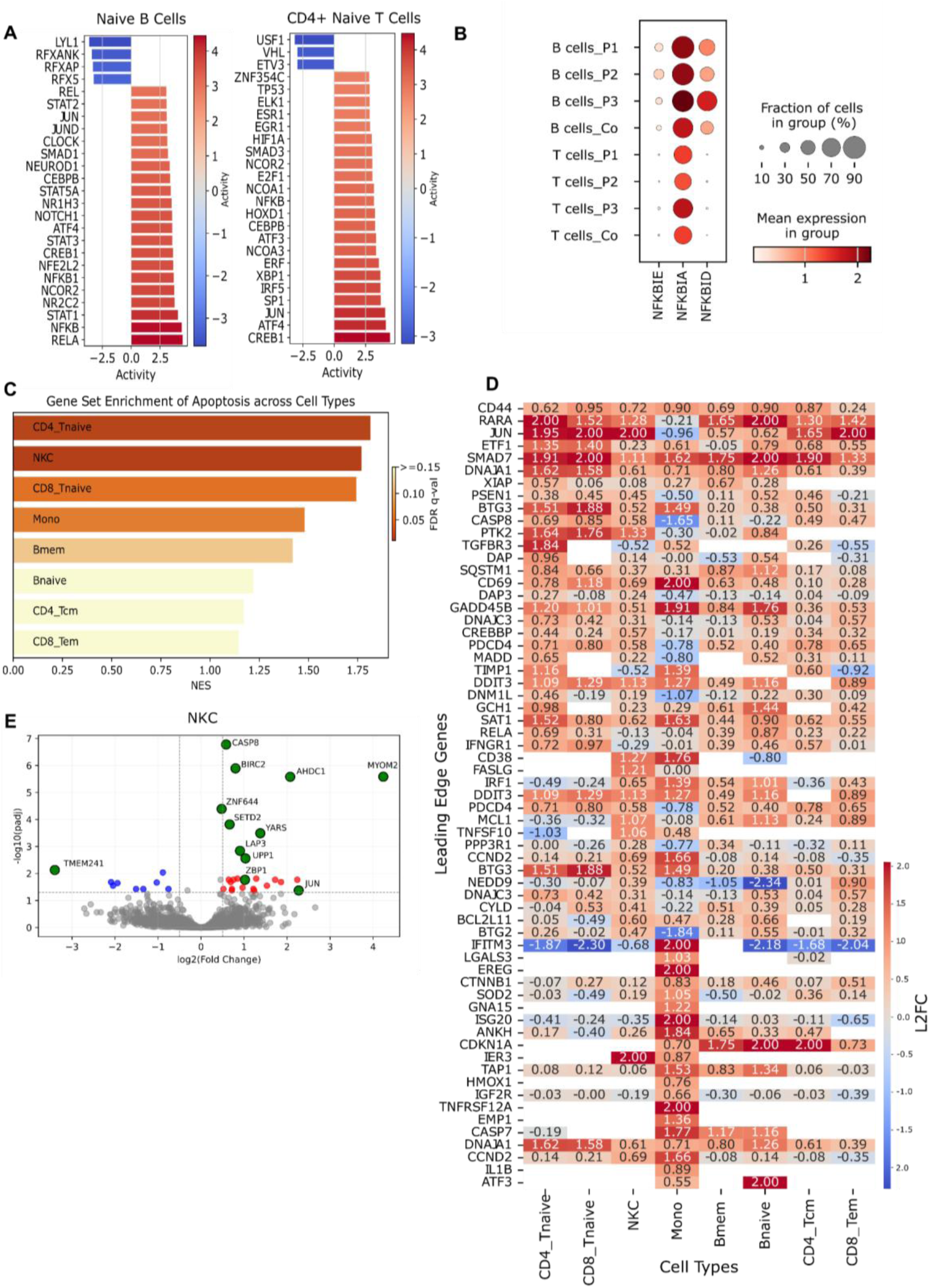
Differential gene expression analysis of single-cell RNA seq data. **(A)** Bar plots reflecting relative transcription factor activity scores based on the expression of known targets of the indicated transcription factors for naïve B cells and naïve CD4^+^ T cells, respectively. **(B)** Dot plot showing the expression if NF-κB inhibitors in patient and control B and T cells. **(C)** Bar plot showing the Gene Set Enrichment Analysis (GSEA) enrichment of the MsigDB Term “Apoptosis” in differentially expressed genes between patient and control cells of the indicated cell types. FDR, False Discovery Rate; NES, Normalized Enrichment Score. **(D)** Heatmap showing the log2 fold-change in expression between patient and control cells for the GSEA “Apoptosis” leading edge genes derived from the analysis in T and NK cells as well as monocytes. **(E)** Volcano plot for the differential expression analysis between patient and control NK cells. The top 10 differential as well as biologically relevant genes are highlighted.

